# Single-cell RNA-sequencing of Herpes simplex virus 1-infected cells identifies NRF2 activation as an antiviral program

**DOI:** 10.1101/566992

**Authors:** Emanuel Wyler, Vedran Franke, Jennifer Menegatti, Kocks Christine, Anastasiya Boltengagen, Samantha Praktiknjo, Barbara Walch-Rückheim, Nikolaus Rajewsky, Friedrich Grässer, Altuna Akalin, Landthaler Markus

**Author notes:** These authors contributed equally to this work (Emanuel Wyler, Vedran Franke).

## Abstract

Herpesvirus infection initiates a range of perturbations in the host cell, which remain poorly understood at the level of individual cells. Here, we quantified the transcrips of single human primary fibroblasts during the first hours of lytic infection with HSV-1. By applying a generalizable analysis scheme, we defined a precise temporal order of early viral gene expression and found unexpected bifurcations and bottlenecks. We identified individual host cell genes and pathways relevant in early infection by combining three different computational approaches: gene and pathway overdispersion analysis, prediction of cell-state transition probabilities as well as future cell states. One transcriptional program, which was turned on in infected cells and correlated with increased resistance to infection, implicated the transcription factor NRF2. Consequently, Bardoxolone methyl, a known NRF2 agonist, impaired virus production, suggesting that NRF2 activation restricts the progression of viral infection. Our study provides novel insights into early stages of HSV-1 infection and serves as a general blueprint for the investigation of heterogenous cell states in virus infection.

## Introduction

Herpes simplex virus 1 (HSV-1) is one of nine known Herpes viruses that affect humans. Although an estimated 80% of the worldwide population is infected in a quiescent, latent form, the virus may cause a variety of diseases during lytic replication and reactivation [1]. A hallmark of HSV-1 is the way it alters the cellular RNA metabolism and RNA content on many levels. On one hand, viral mechanisms activate transcription of viral genes and interfere with splicing regulation, RNA polymerase II transcription, and RNA stability to inhibit synthesis of cellular proteins [2-8]. On the other hand, virus entry affects a range of cellular pathways, which may in turn lead to transcriptional activation or repression of downstream target genes [9].

Viral infection is a dynamic process driven by the interplay of anti-viral cellular pathways and viral mechanisms which evolved to suppress them. Incoming HSV-1 virions bind to receptors on the cell surface, including NECTIN1/NECTIN2, the TNF receptor superfamily member 14 (TNFRSF14) [10] and the members of the integrin family [11], resulting in activation of cellular pathways such as NF-κB, or of the transcription factors IRF3 and IRF7 [11]. After entry into the host cell, a range of pattern recognition receptors sense viral DNA or RNA, such as the cyclic GMP-AMP synthase MB21D1 (also known as cGAS) [12, 13]. Eventually, these pathways can lead to the induction of inflammatory cytokines and interferons [10].

The characterization of cellular heterogeneity due to activation of different host pathways and the progression of viral infection is of great interest. Recent single-cell RNA-sequencing (scRNA-seq) efforts provided an unbiased characterization of virus–host interactions in individual cells which are masked at the population level [14-22]. However, deeper insights into unique molecular signatures and discovery of specific cell subsets could be obtained by increasing sequencing depth and the application of advanced analytical approaches to study viral infections.

Here, we profiled deep transcriptomes of tens of thousands of individual cells harvested before and at several times post HSV-1 infection. Our results related the progression of infection to cell cycle phases, and defined a precise temporal order of viral gene expression. The depth of data and using unspliced mRNA as a predictor for future cell states allowed us to connect the course of infection to the activity of specific host cell genes and pathways. Particularly, we investigated the relationship of HSV-1 infection and the transcription factor NRF2, which is activated during infection, and demonstrated that the NRF2 agonist Bardoxolone methyl impairs a productive viral replication. Overall, our study provides novel insights into early stages of HSV-1 infection, and provides an analytical framework to study viral infections using scRNA-seq.

## Results

### Single-cell RNA-sequencing of HSV-1 infected primary human fibroblasts

To investigate the heterogeneity of molecular phenotypes in the first hours of viral infection, we infected primary human fibroblasts (NHDF) with HSV-1 at an MOI of 10 (Fig. 1a,b) and profiled the transcriptomes of uninfected cells as well as cells harvested at 1h, 3h and 5h post infection using the droplet-based single-cell sequencing (Drop-seq) [23, 24]. For further analysis, only cells with more than 2000 detected genes were used, a threshold that has been previously shown to reduce technical variability [25]. An overview of the dataset (Table S1), number of characterized cells (Table S2), distribution of unique molecular identifiers (nUMI) and number of detected genes (nGene) (Supplementary Fig. 1a), as well as correlation between scRNA-seq and bulk RNA-seq (Fig. S1B) are provided in Supplemental data. Low-reproducibility genes (Table S3) were subsequently omitted or flagged.

**Figure 1.**
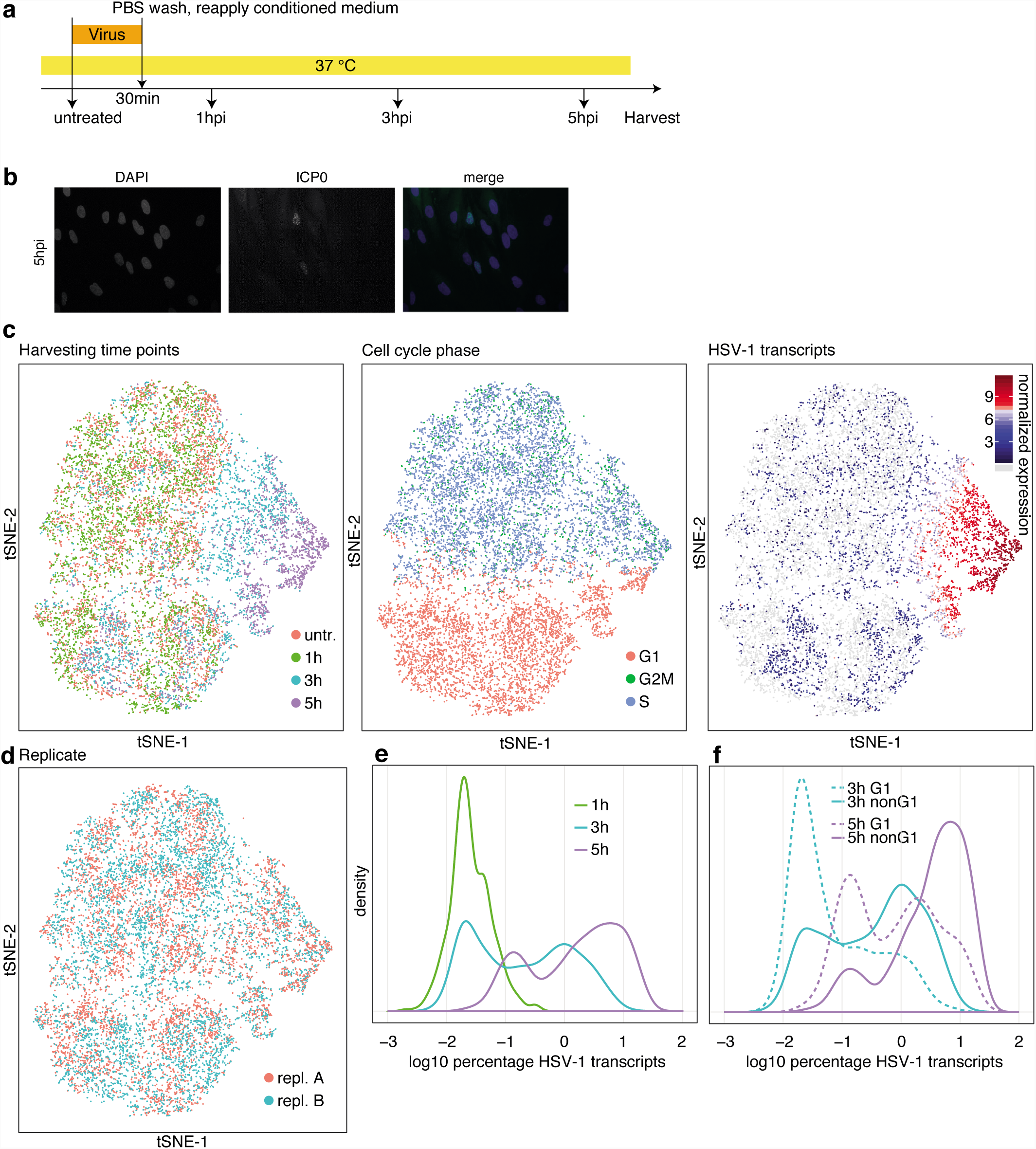
Investigating HSV-1 infection using single-cell RNA-sequencing of primary human fibroblasts shows cell cycle dependency. a, Infection protocol. In parallel to single-cell RNA-sequencing, cells were harvested for bulk mRNA-sequencing and ICP0 immunofluorescence staining. b, ICP0 immunofluorescence staining at 5hpi. c, Global display of scRNA-seq data as tSNE maps. Cells were colored by, from left to right, harvesting time points, cell cycle phase, and the normalized values of the sum of HSV-1 transcripts as a marker for the progression of infection. Cells without HSV-1 transcripts are in light gray. d, tSNE maps with cells colored by replicate. e, Relative densities of the percentage of viral transcripts (log10 transformed) per cell for the three time points post infection. f, Relative densities of the percentage of viral transcripts per cell (log10 transformed) for G1 and non-G1 cells for cells harvested at 3hpi and 5hpi.

The analyzed cells clustered based on harvesting time point, cell cycle markers, and the amount of viral mRNA, suggesting that the strongest contributors to cellular variability were cell cycle state and the progression of infection (Fig. 1c). However, cells did not group by biological replicates, indicating that replicates provided comparable and reproducible data (Fig 1d).

The distribution of the viral gene expression per single cell at the different harvesting time points indicated the progression of infection over time. (Fig. 1d). Separating cells based on their cell cycle state (G1 vs. non-G1) showed that, for a given harvesting time point, non-G1 cells generally contain more viral transcripts (Fig. 1e), suggesting that S, G2 and M phase cells are more susceptible to viral infection, and/or that the infection progresses faster in these cells.

Consequently, at 5 hpi we observed that cells bearing high levels of HSV-1 mRNA (8-30%) showed a lower nUMI count when related to the number of detected genes (Supplementary Fig. 1c), indicating less complex transcriptomes due to large number of viral transcripts and/or reduction of fibroblast encoded mRNAs likely as a consequence of the host cell shut-off [2].

### Precise definition of the temporal order of viral gene expression

Within lytic infection, viral genes have been classified as immediate early (alpha), early (beta), and late (gamma1, gamma2) [26, 27]. Since the temporal order of genes expressed from the virus genome is intrinsically encoded in their single-cell expression profiles, we developed an approach to derive a precise model of the early viral gene expression cascade.

As a proxy of the sequential appearance of virus-encoded transcripts, we counted, for each gene, the percentage of infected cells in which it was detected (Fig. 2a). In order to focus on early viral transcriptional events, only cells harvested at 1hpi and 3hpi were used (cells harvested at 5hpi are depicted in Supplementary Fig. 2b). Interestingly, viral gene transcription appears to start mostly around the internal repeats, and around gene UL23 (Fig. 2a).

**Figure 2.**
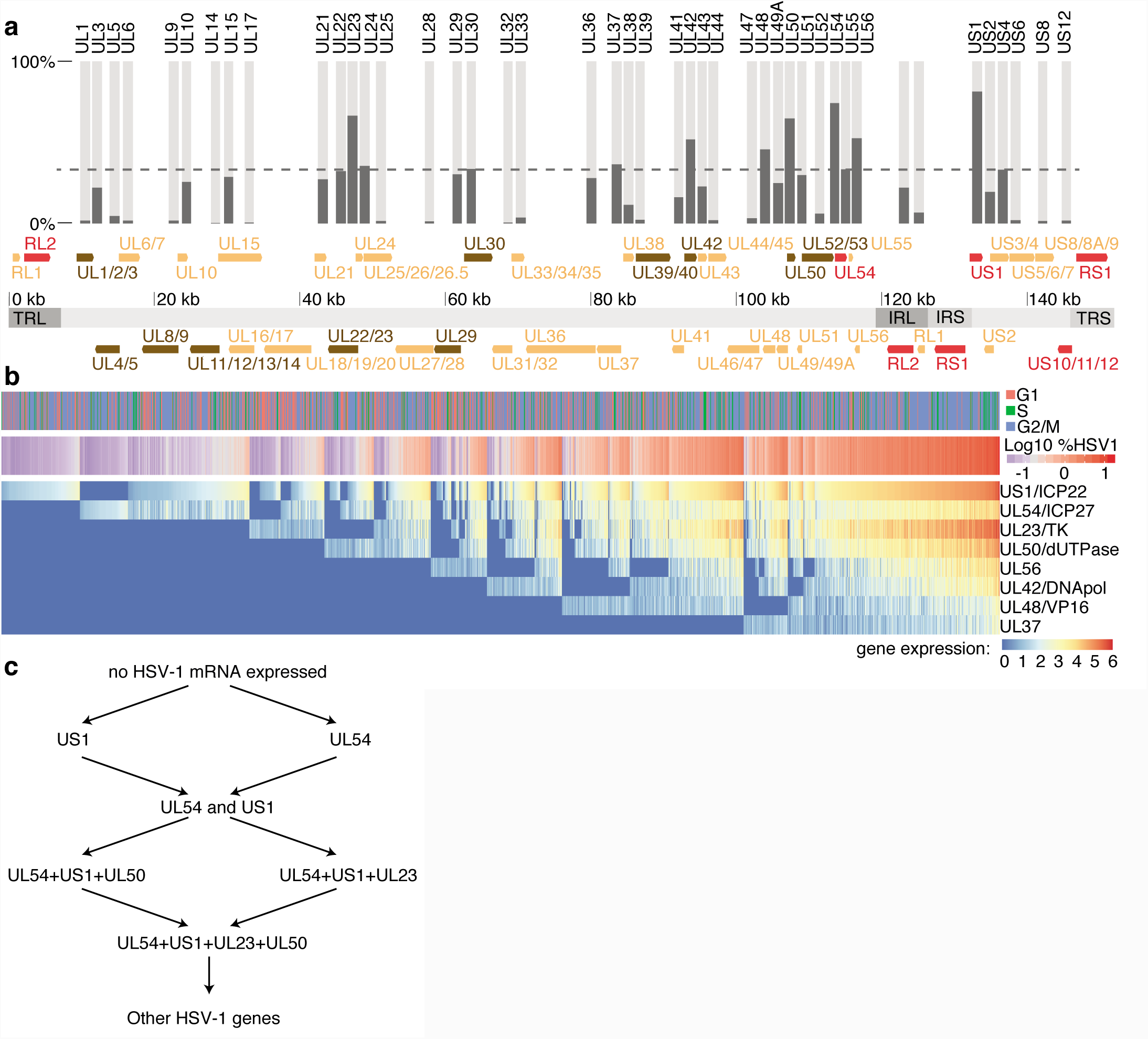
Onset of viral gene expression shows bifurcations and bottlenecks. a, Arrangement of viral genes on the HSV-1 genome. Immediate early genes were shown in red, early genes in brown, and late genes in yellow. Genome segments were shown as light gray (unique regions) and dark gray areas (IRL, IRS: large and small internal repeat region; TRL, TRS: large and small terminal repeat region). For overlapping genes (e.g. UL1, UL2 and UL3), the gene regions were merged. The bar plot on top shows in which percentage of “HSV-1 high” cells (see Fig. S2A for definition) the respective transcript was detected. b, Heatmap of expression values of the first eight expressed viral genes for experiment two with only “HSV-1 high” cells as defined in the main text and Fig. S3A, harvested at 1hpi and 3hpi. Rows (genes) and columns (cells) were sorted as described in the main text. Above the heatmap, cell cycle and log10 transformed percentage of viral transcripts are shown. For the latter, blue colors indicate values in the first peak and second peak of the bimodal distribution from Fig. 1e, respectively. c, Simplified scheme of proposed cell states in early viral infection.

For the first eight viral genes as defined by this ordering, normalized expression values were represented in Fig. 2b. Cells were first sorted according to appearance of the eight genes, and then by the abundance of US1 (coding for ICP22), or, if absent, UL54 (ICP27). Under the assumption that the progression of infection, at least in an early phase, is correlating with the accumulation of viral transcripts, this represents a pseudo-time course of the lytic infection. In agreement with previous findings [26, 27], US1 and UL54 were the first genes to appear.

Early in infection, there are likely three distinct sets of cells, with either US1 or UL54, or both genes expressed. A comparably high number of cells in these three first states suggested that viral transcription progresses slowly until a number of US1 and UL54 transcripts were present, indicating that the two earliest genes US1 and UL54 were transcribed before the UL23/UL50 transcripts are expressed.

As shown above (Fig. 2a), we observed a bimodal distribution of the relative densities of HSV-1 transcript amounts per single cell. We connected the bimodal distribution of viral transcripts per single cell (Fig. 1e) to the sequential expression of viral genes in two ways. First, cells in the first peak bear mainly transcripts from one or two viral transcripts (Fig. 2b, Supplementary Fig. 2b). Second we generated smoothened two-dimensional densities of cells with plotting the number of detected genes vs. the percentage of viral transcript per cell (Supplementary Fig. 2c). Together, this indicates that the bimodality in Fig. 1e arises from the transition between US1/UL54 expression and later stages of infection.

Calculating the likelihood of transitions between single cells based on diffusion maps [28, 29] indicated “bottlenecks” in the accumulation of HSV-1 transcripts, as represented by clusters of high transition probabilities (Supplementary Fig. 2d). Cells in the two-gene state (US1 and UL54) showed a higher likelihood to transition into the four-gene state, deriving a early gene expression cascade shown in Fig. 2c.

### The small GTPases RASD1 and RRAD are expressed in infected cells and reduce virion production

To study changes in viral gene expression upon infection, we first examined differential gene expression by bulk mRNA sequencing (Supplementary Fig. 3a). To identify genes differentially expressed only in infected cells, we plotted the correlation of host gene expression to viral transcripts in single cells against the maximal fold change in the population mRNA-seq data (Fig. 3a).

**Figure 3.**
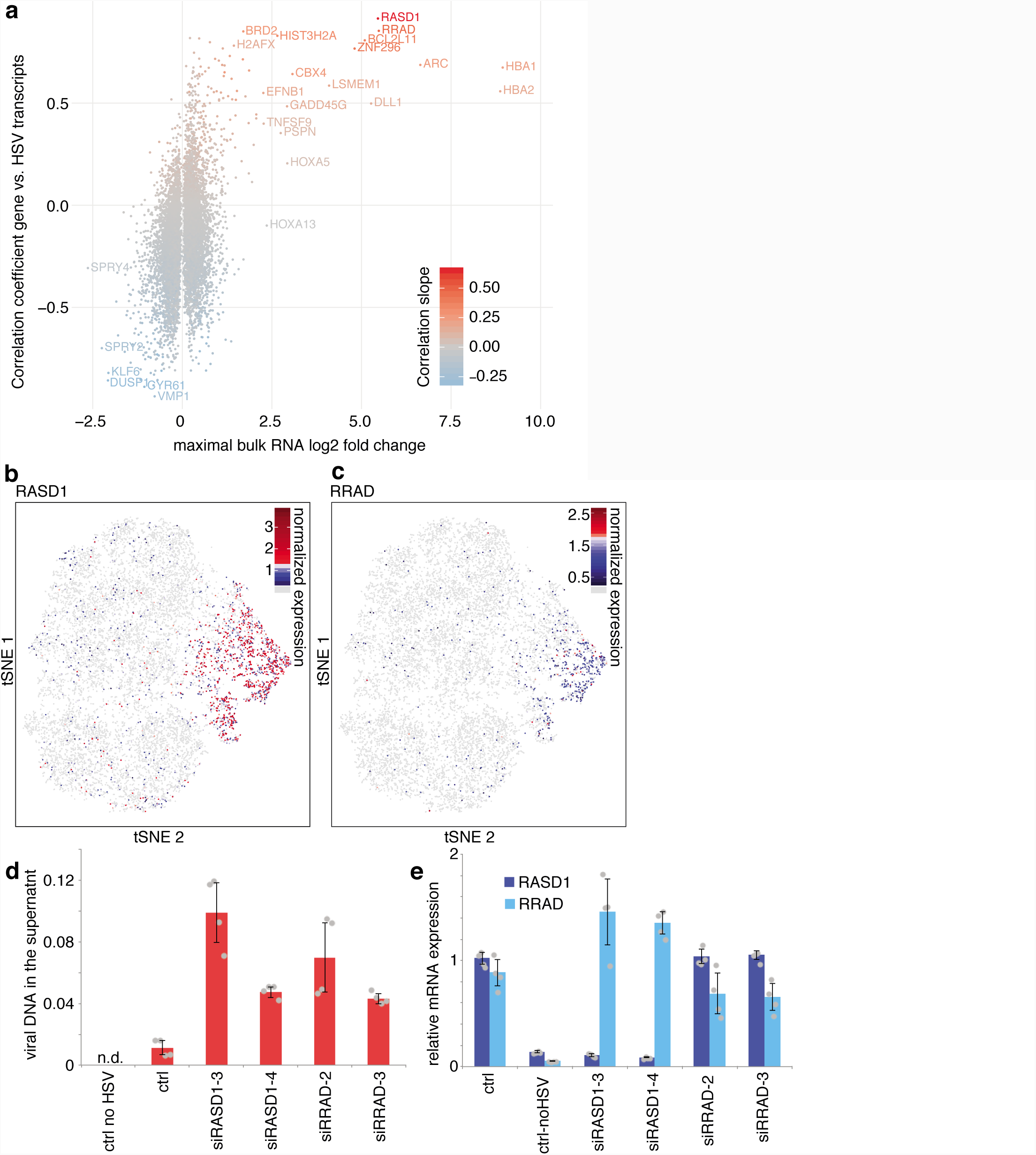
Relating host cell gene expression to viral transcription reveals candidates for modulation of infection. a, Relationship between differentially expressed genes and viral transcription. The horizontal axis shows the maximal log(2) transformed bulk RNA-seq fold change of the three time points after infection compared to uninfected cells, the vertical axis the linear regression coefficient of the gene expression with the sum of viral transcripts in bins of 20 cells, using only high HSV-1 cells as defined in Supplementary Fig. 2a. Color represents the slope of the linear regression. bc, Distribution of normalized expression of RASD1 (b) and RRAD (c) on the tSNE projection introduced in Figure 1. Cells without detectable expression are colored in light gray. d, RNAi of HSV-1 transcription dependent factors RASD1 and RRAD. Viral DNA in the cell culture supernatant was measured using qPCR and quantified using serial dilutions of a virus stock with known activity (a value of 1 corresponding to 10^6 PFU/ml). Barplots indicate means, error bars denote standard deviations, the individual measurement values are shown as grey dots. e, RT-qPCR of RNAi samples. RASD1 and RRAD mRNA levels were normalized using GAPDH mRNA values and to non-infected control cells. The individual measurement values are shown as grey dots.

Among host genes, activated only in infected cells, we identified hemoglobin alpha genes, HBA1 and HBA2 (Fig. 3a), which were previously shown to be induced by the viral transcription factor ICP4 [7]. Additionally, two genes encoding Ras-related small GTPases, RASD1 (Fig. 3b) and RRAD (Fig. 3c), showed strong positive correlations with viral transcripts. Similarly, we found that RASD1 and RRAD were up-regulated in previously published microarray, RNA-seq, and ribosome profiling datasets [5, 6, 30, 31], indicating that the induction of these two genes is independent of the cell type and virus strain. To investigate the influence of RASD1 and RRAD on viral production, we depleted RASD1 and RRAD by siRNA knockdown prior to infection (Fig. 3d,e), and observed increased viral DNA in the supernatant 24 hpi, suggesting that both genes provide some antiviral activity.

### Distinct subpopulation of cells marked by mutually exclusive NQO1 and SULF1 gene expression

To identify distinct subpopulations of cells, defined by a particularly high or low expression of specific marker genes, we used an overdispersion analysis [32]. Not surprisingly, cell cycle, viral genes and host cell genes correlating with viral infection appeared as the strongest markers (Fig. 4a-e). In addition, two other gene sets, not directly related to the progression of infection and the cell cycle, outlined distinct subpopulations. Importantly, the genes discussed here do not necessarily by themselves influence the infection, but rather are indicators of a specific cellular state that could favor or impair the infection.

**Figure 4.**
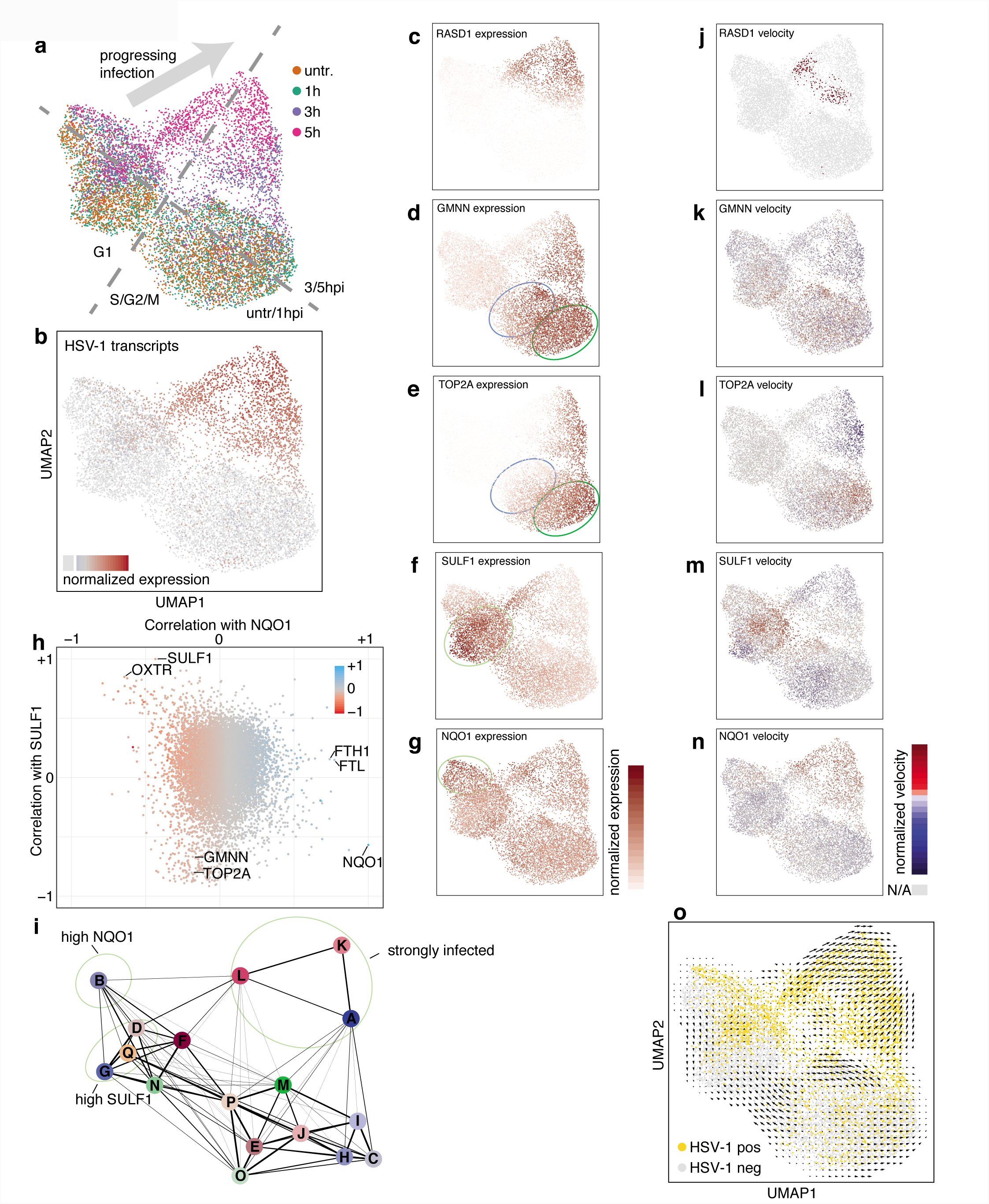
Overdispersed genes mark subpopulations of cells with distinct transition probabilities into infection. a, Cells were projected on a two-dimensional map using UMAP according to gene expression values, with cells colored by harvesting time point. Main distinguishing features were marked in grey. b, Cells colored by amount of HSV-1 transcripts. Cells without HSV-1 transcripts are colored in light gray. c-g, Cells colored by expression values of the indicated genes. Color scales are shown to the right of these panels. areas containing cells with relatively high expression levels were marked with a light green ellipse. The dark green and blue ellipse denote S and G2/M phase cells, respectively. h, correlation of gene expression with NQO1 expression (horizontal axis) and SULF1 expression (vertical axis) in cells harvested at 3hpi. Every dot represents a gene, colored by the slope of the linear correlation with NQO1. Selected genes were labeled by name. i, Groups of cells according to PAGA were labeled A to Q. The line thickness of the connections (graph edges) between the cell clusters (graph nodes) indicate the likelihood that cells can move from one cluster to the other (right). Areas of interest are marked with light green ellipses. j-n, Cells colored by RNA velocity values of the indicated genes. Cells for which no value could be calculated were colored in light gray. o, RNA velocity arrows projected on the UMAP. Cells containing at least one detected viral transcript were colored in light orange, all others in light gray.

The first subpopulation was marked by high mRNA levels of the sulfatase SULF1 (Fig. 4f), and the oxytocin receptor OXTR (Supplementary Fig. 4a). The second set was characterized by high mRNA levels of the NAD(P)H quinone dehydrogenase 1 NQO1 (Fig. 4g), the Ferritin heavy chain 1 FTH1 (Supplementary Fig. 4b), as well as the Ferritin light chain FTL (not shown) and Sequestosome 1 SQSTM1 (not shown).

Whereas SULF1 was previously shown to be induced by TNFα in MRC-5 fibroblasts [33], we found no clear activators for OXTR, but indications that proinflammatory cytokines such as IL-1β, IL-6, and TNFα might induce its transcription [34]. For the activation of NQO1 and the correlating genes FTH1, FTL, and SQSTM1, several reports pointed to the transcription factor NFE2L2 (also known as NRF2) [35-37].

Pairwise gene expression analysis revealed a strong anticorrelation of SULF1 and NQO1 expression in single cells, whereas FTH1 and FTL correlated well with NQO1 (Fig. 4h). On the other hand, OXTR expression levels correlated with SULF1 but not with NQO1, indicating the existence of subpopulations of cells with distinct marker gene expression.

Next we related these subpopulations to the progression of infection. To this end, we used graph abstraction (PAGA) [38], which calculates transition probabilities between groups of cells (Fig. 4i). Cells with high NQO1 levels, and therefore high preceding NRF2 activity, had a relatively low probability to progress into the infection, compared to cells from groups D and F.

### RNA velocity shows interruption of the cell cycle by HSV-1 infection

To infer expression dynamics in individual cells, we used RNA velocity [39]. RNA velocity uses the ratio of spliced to unspliced mRNAs to derive transcriptional rates. RNA velocity values are shown for the genes mentioned above (Fig. 4j-n). Cells in the G1 phase that show high GMNN RNA velocity (Fig. 4k), are therefore cells which are about to enter the S phase. Interestingly, cells in a more progressive state of infection had low RNA velocity values for both GMNN and TOP2A, (Fig. 4k, l, top part of the maps), which reflects the interruption of the cell cycle by the virus [40]. In addition, RNA velocity allows the prediction of the future state of individual cells on a timescale of hours. The progression of infection clearly emerged as the predominant transition (Fig. 4o), confirming the validity of the approach. Since viral genes barely have introns, the directionality of infection progression is driven by virus-induced host cell genes such as RASD1 (Fig. 4c,j).

### The transcription factor NRF2 is activated upon viral infection

We used the inferred RNA velocity (transcriptional rates) as a precise read out of activation of upstream signaling pathways. To visualize pathway activity, we mapped the cells on a two-dimensional embedding not based on the mature mRNA expression levels as in Fig. 4, but based on the RNA velocity values (Fig. 5a,b). Cells with high SULF1 transcription (Fig. 5c, lower panel), which could reflect TNFα-responsive cells, clustered together in the lower left part. These cells are mostly harvested at later time points but do not show high expression of HSV-1 transcripts, reflecting the established antiviral activity of TNFα [41, 42]. Cells with larger amounts of viral transcripts showed relatively high NRF2 activity as deduced from high NQO1 RNA velocity (Fig. 5d, lower panel) suggesting that NRF2 is activated as a part of the cellular defense against the progressing infection.

**Figure 5.**
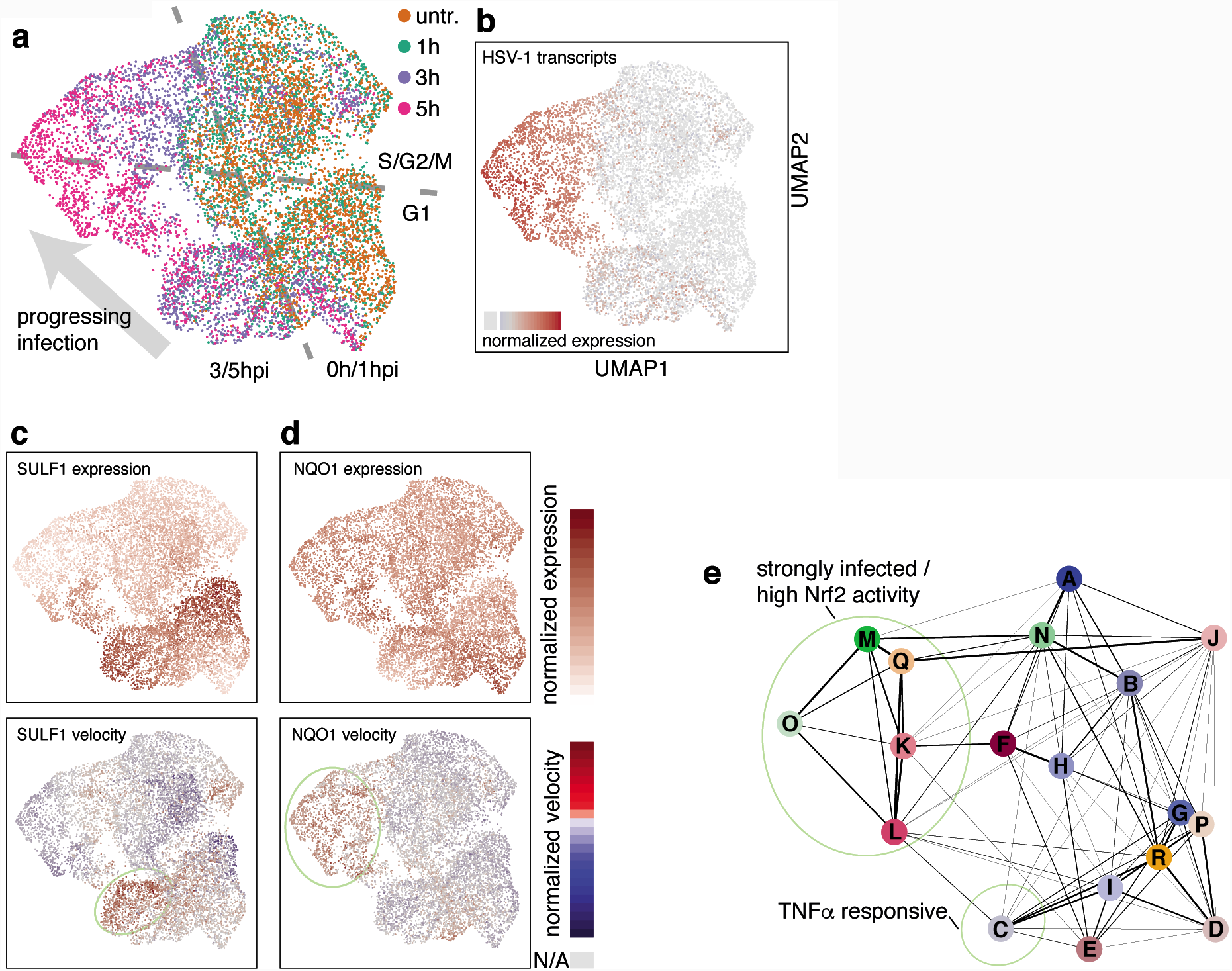
Clustering of cells by RNA velocity reveals subpopulations with common transcriptional activity. a, Cells were projected on a two-dimensional map using UMAP according to RNA velocity values, with cells colored by harvesting time point. Main distinguishing features were marked in grey. b, Coloring by amount of HSV-1 transcripts. c and d, Cells colored by expression values (top panels) and RNA velocity values (bottom panels) of the indicated genes. Color scales are shown to the right of these panels. For expression values, areas containing cells with relatively high expression levels were marked with a light green ellipse. For RNA velocity, cells for which no value could be calculated were colored in gray. e, Groups of cells according to PAGA were labeled A to R. Transition probabilities between clusters are proportional to the line thickness between the clusters (right).

We again applied the PAGA algorithm to cells clustered based on transcriptional activity (Fig. 5e). The highest probability to proceed into infection are cells in the S/G2/M phases (groups F, N, Ja, supporting again that these phases of the cell cycles favor infection. On the other side, cells with high SULF1 transcription are in cluster C, which shows a comparably lower probability to transition into infection, confirming the lower susceptibility of cells with an activated TNFα response to HSV-1.

To summarize, we made two observations regarding NRF2 activity and HSV-1 infection. The analysis of mature mRNA distribution showed that cells with a high level of transcripts of NRF2-driven genes, and therefore high preceding NRF2 activity, have a low transition probability into later stages of infection (Fig. 4). Looking at RNA transcription however showed that cells at later stages of infection appear to respond by increasing transcription of NRF2 target genes (Fig. 5), which could reflect a cellular defense mechanism against HSV-1 infection.

In order to strengthen our interpretation of the data, we also analyzed an independent experiment with again two biological replicates, where a procedure for synchronized infection at 4 °C [43] was applied (Supplementary Fig. 5a, Supplementary Table 1). Whereas overall the infection progressed less (compare Supplementary Fig. 5b with Fig. 1e), key aspects of our analysis were reproducible. This included the cell cycle dependency (Supplementary Fig. 5c), the viral gene expression cascade (Supplementary Fig. 5d) and the induction of RASD1/RRAD (Supplementary Fig. 5e). Due to the weaker infection, there are less infected cells as a separated part on the two-dimensional projection (compare Supplementary Fig. 5f,g with Fig. 4a,b). Still, the anti-correlation of NQO1/SULF1 and the low transition probability of cells with high NQO1 levels into infection was observed (Supplementary Fig. 5f-j).

### The NRF2 agonist Bardoxolone methyl restricts HSV-1 infection

Our analysis suggested an activation of NRF2 in a subset of infected cells. Under physiological conditions, NRF2 is repressed by KEAP1, which sequesters NRF2 and facilitates its ubiquitination and degradation [44]. Upon disruption of this interaction in response to oxidative stress, NRF2 translocates into the nucleus, and induces transcription of a number of target genes, including NQO1. Using the NRF2 agonist Bardoxolone methyl [45] at final concentrations of 0.1-0.4 μM, we observed an increase in NQO1 mRNA expression in primary fibroblasts (Fig. 6a) and in HEK 293 cells (Supplementary Fig. 6a), as expected for NRF2 activation [46, 47]. At the concentrations applied here, Bardoxolone methyl did not have any apparent effect on cell growth (data not shown). Since, as described above, the progression of infection depends on the cell cycle, we also probed mRNA levels for the cell cycle markers GMNN and TOP2A introduced in the previous sections (Supplementary Fig. 6d). In the primary fibroblasts but not HEK 293 cells, the levels of these mRNAs were somewhat reduced, indicating that Bardoxolone methyl could also dampen cell cycle progression or promote cell cycle exit.

**Figure 6.**
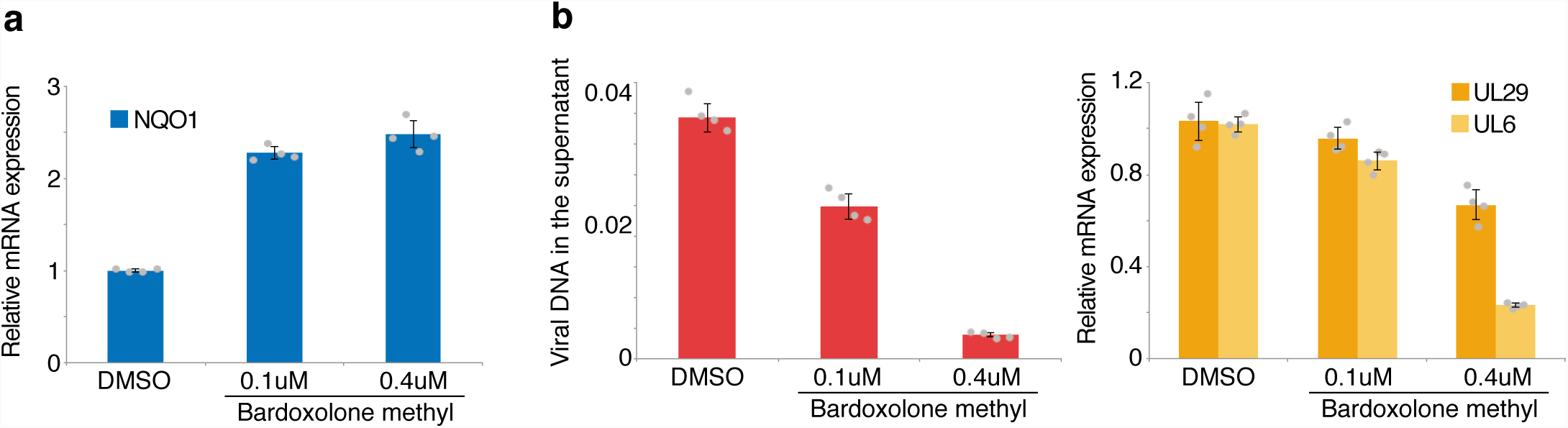
Bardoxolone methyl impairs HSV-1 infection. a, primary human fibroblasts (NHDF) were treated for 16h with solvent (DMSO) or different concentrations of Bardoxolone methyl, followed by RNA isolation. NQO1 mRNA levels were probed using RT-qPCR. b, NHDF cells were infected with HSV-1 at an MOI of 1. After removal of virus inoculum, washing with PBS, and adding back conditioned mediums, solvent or different concentrations of Bardoxolone methyl were added. At 16hpi, viral DNA was probed in the supernatant using qPCR (left panel) and viral mRNAs in the RNA isolated from the cells using RT-qPCR (right panel). For all panels, barplots indicate means, error bars denote standard deviations, the individual measurement values are shown as grey dots.

Next, we tested whether Bardoxolone methyl modulates HSV-1 infection. To this end, cells were infected and Bardoxolone methyl was added immediately after removal of the virus inoculum. At 16hpi, Bardoxolone methyl treatment reduced the levels of produced virions, as measured by the amount of viral DNA in the supernatant (Fig. 6b, left panel, and Supplementary Fig. 6b, left panel). The mRNA of the early UL29 and particularly the late UL6 gene (Fig. 6b, right panel, and Supplementary Fig. 6b, right panel) was also reduced, suggesting that late stages of viral transcription/replication and/or virion production are impaired in cells treated with Bardoxolone methyl. During infection, this treatment led to an increase in NQO1 mRNA levels (Supplementary Fig. 6c).

## Discussion

We performed single-cell RNA-sequencing of primary human fibroblasts at early stages of infection with HSV-1. Using the resolution of scRNA-seq, we showed that cells in the S/G2/M cell cycle phases bear more viral transcripts. The relationship between the cell cycle and the progression of infection has been studied for decades [48-51], with most reports pointing to a G1 or G1/S arrest upon HSV-1 infection. Single-cell RNA sequencing now allows for perturbation-free analysis of the relationship between the cell cycle and progression of infection, reducing the risk of experimental artifacts. We have seen that cells in S/G2/M phases on average bear more viral transcripts (Fig. 1f). This is corroborated by recent findings that the activity of CCNE1 (also known as cyclin E) and CDK2, which promote the G1/S transition, correlates with productive infection [52]. Similarly, high levels of the G1/S transition marker GMNN (also known as geminin) favored infection in microscopy-based experiments [53]. We observed that, under the conditions used here, the fibroblast populations doubled every 40-50 hours. Since about half of the measured cells were in S/G2/M phases, these would last more than 20 hours, much longer than the intervals between infection and harvesting. Our results therefore indicate that cells in S/G2/M phases provide a more favorable environment to establish the infection compared to G1 cells. Extending measurements to later time points could then also detail the cell cycle state in which infected cells are arrested. Herpes viral genes are classified into immediate early, early, and late [1]. Still, studies so far averaged large numbers of cells, and were not able to distinguish how viral gene expression starts in individual cells. scRNA-seq allowed us to find two bifurcations at the onset of viral gene expression, and one or several “bottlenecks” in the viral gene expression cascade (Fig. 2). Investigating viral gene expression patterns from sufficiently large number of cells (Fig. 2) combined with diffusion maps (Fig. S2d) provides a methodological framework that could be used to investigate other cases of pathogen gene expression.

Virus infection leads to changes in host cell transcription. We have identified a number of deregulated genes in bulk-RNA sequencing, but only using the scRNA-seq data we could distinguish on how the deregulation is related progression of infection in the same cell. Two small GTPases, RASD1 and RRAD, were directly induced by the infection. We have observed that both possesses antiviral properties (Fig. 3). Interestingly, these two genes are usually not expressed in epithelial cells. They are however putative targets of the primate specific gene DUX4 [54]. This germline transcription factor was recently shown to be induced in HSV-1 infected cells [55]. Correlating host gene expression with viral gene expression generally emerges as a focus in scRNA-seq studies of virus infected cells [15, 17], and comparative studies might reveal common topics and modulators of viral infection.

The emergence of pseudo-time inference algorithms for scRNA-seq data [56], enabled us to order cells along defined trajectories and made it possible to deduce relationships between biological mechanisms, without artefact-prone perturbances such as gene knockdowns/knockouts. We first used over-dispersion analysis [32] to find genes with inhomogeneous expression across the entire dataset, which might therefore play a role in the infection (Fig. 4). The relationship of these genes to the progression of infection was then analyzed using RNA velocity [39] and graph abstraction [38]. We focused on two groups of genes, one represented by NQO1 (target of the transcription factors NRF2), and the other by SULF1 (associated with the TNFα response). Importantly, in bulk RNA-seq data, these genes do not appear as differentially expressed and would likely be omitted from subsequent studies.

NRF2 has previously been linked virus infections [57], however without a clearly defined role. Particularly, in a recent Influenza virus scRNA-seq study, NRF2 activity has been associated with high amounts of viral transcripts [15]. For Rotavirus, a dsRNA virus, NRF2 activation was recently shown to reduce virus production [58]. Our analysis suggests that NRF2 activity in a subset of cells leads to an “escape state”, diverging from a progressing lytic infection (Fig. 4), and that NRF2 becomes active upon infection (Fig. 5). For Herpes infections, reports on the role of NRF2 so far are mixed. Treatment of mice with the NRF2 agonist tert-butylhydroquinone protected them from MCMV infection [59]. In contrast, HSV-1 infection in A549 was drastically reduced upon a two-day RNA interference of NRF2 [60]. This treatment however induced the antiviral genes STING and IFI16, which could confound the direct effect of the NRF2 depletion. In another study, as observed here for HSV-1, KSHV infection induced NRF2 [61]. We followed up on our observations by using the NRF2 agonist Bardoxolone methyl, which is currently in phase 3 clinical trials for treating chronic kidney disease [62, 63]. Interestingly, Bardoxolone methyl impaired the infection. At which stage of the infection viral production is impaired, and whether by mere presence of nuclear Nrf2 or via its target genes, as well as the relevance of reported NRF2 interaction with the NF-kappaB pathway, remains to be investigated.

In summary, our study provides a detailed analysis of the events at the beginning of HSV-1 infection in primary human fibroblasts. We could relate the activity of single genes to the progression of infection. Using overdispersion and cellular trajectory analysis allowed the identification of biologically relevant pathways, demonstrating how scRNA-seq can provide an unprecedented insight in the heterogeneity in viral infections.

## Acknowledgments

The authors wish to thank Julia Madela, Sebastian Voigt, Lars Dölken, Benedikt Kaufer and Jermaine Voß and members of the Landthaler and Rajewsky labs for fruitful discussions and support.

## Author contributions

E.W. performed experiments, analyzed data and wrote the manuscript, V.F. performed coding and data analysis and contributed to manuscript writing, J.M. did cell culture and virus infection for scRNA-seq, bulk RNA-seq, and RNAi experiments, C.K. contributed to manuscript writing and performed DropSeq experiments together with A.B., S.P. contributed to manuscript writing and data analysis, B.W.-R., N.R., and F.G. supervised parts of the project, A.A. and M.L. were responsible for overall supervision.

## Data availability

Raw sequencing reads as well as raw read counts for both the bulk RNA-seq and the single-cell RNA-seq, along with normalized counts and tSNE coordinates and cell cycle information, are available in the NCBI GEO repository, accession number GSE123782.

## Code availability

Custom R, Python, Perl, awk and bash scripts are readily available from the authors upon request.

## Supplementary Tables

Supplementary Table 1: sample overview

Supplementary Table 2: variable genes

Supplementary Table 3: numbers of quantified cells, cells with HSV-1 transcripts

Supplementary Table 4: qPCR primers

## Supplementary Figure Legends

Figure S1 | a, Violin plots showing the distribution of number of unique molecular identifiers and number of detected genes in the individual scRNA-seq samples. b, Correlation of bulk RNA-seq with scRNA-seq data. Shown are scatter plots (lower left part) and linear correlation coefficients (upper right part) of bulk polyA RNA-seq values (horizontal axes, log(2) transformed fpkm values) and scRNA-seq values (vertical axes, log(2) transformed sums of scRNA-seq counts) of merged replicates, c, Cells harvested at 5hpi containing at least one viral transcript were binned according to the percentage of viral transcripts per cell, and the log2 transformed nUMI/nGene values plotted as smoothed means with 95% confidence intervals in gray. Percentage ranges of the bins are indicated below the plot.

Figure S2 | a, Relationship between normalized levels of HSV-1 transcripts and number of detected genes shows two categories of cells. Cells with at least one viral transcript were sorted by normalized HSV-1 transcript UMIs. For each cell, the red dot represents the log(2) transformed sum of HSV-1 transcript UMIs (left axis), and the green dot the number of genes detected (right axis). Note that only cells with more than 2000 detected genes were used, thus the lower apparent limit for the green dots. Light and dark gray bars designate “low HSV-1” cells and “high HSV-1” cells according to the bimodality of the plot. The horizontal gray line denotes the cutoff expression between high and low. For characterization of the gene expression cascade of viral transcripts, only the 3896 “high HSV-1” cells were used, in order to reduce the sampling error caused by the detection rate. b, Heatmap of expression values of the first eight expressed viral genes for experiment two with only “HSV-1 high” 5hpi cells as defined in the main text. Rows (genes) and columns (cells) were sorted as described in the main text for Figure 3b. Above the heatmap, cell cycle and log10 transformed percentage of viral transcripts are shown. c, Smoothened two-dimensional densities of the number of detected viral genes (vertical axis) in relationship to the percentage of viral transcripts (horizontal axis) for the set of cells used for Fig. 2b. Relative densities as in Fig. 2b are shown on top. d, Clustered heatmap of transition probabilities on the diffusion maps of cells bearing HSV-1 transcripts. Left part, clustered probabilities colored from white (low) to red (high). Columns and rows represent cells in the same order. Right part, expression of the first eight viral genes. The dotted rounded rectangle represents the cluster of transition probabilities between the states denoted under the heatmap. For this analysis, only S/G2/M phase cells were used, since the difference between G1 and non-G1 single-cell transcriptomes otherwise dominates the transition probabilites. Also, cells harvested at 5hpi are left out in order to better study early viral gene expression.

Figure S3 | a, Differential bulk mRNA expression comparing expression from 1hpi (left panel), 3hpi (middle panel), and 5hpi (right panel) with untreated cells. Outliers were marked by name. Horizontal axis denote log(2) transformed fold-changes, vertical axis log(10) transformed p-values. The color represents the log(2) transformed counts per millions over all time points. Low-reproducibility genes (see Supplementary Table S3) are displayed as light gray dots.

Figure S4 | ab, cells colored by expression values (top panels) and RNA velocity values (bottom panels) of the indicated genes. Color scales are shown to the right of these panels. For expression values, areas containing cells with relatively high expression levels were marked with a light green ellipse. For RNA velocity, cells for which no value could be calculated were colored in light gray. c, Cells harvested at 3hpi were sorted by normalized SULF1 transcript UMIs. For each cell, the red dot represents the log(2) transformed amount of SULF1 transcript UMIs (left axis), and the green dot the number of genes detected (right axis). Note that only cells with more than 2000 detected genes were used, thus the lower apparent limit for the green dots. Light and dark gray bars designate low SULF1 cells and high SULF1 cells according to the bimodality of the plot. The horizontal gray line denotes the cutoff expression between high and low. Only high SULF1 cells were used to calculate the correlations in Fig. 4h.

Figure S5 | a, Infection protocol for the synchronized infection at 4 °C. In parallel to single-cell RNA-sequencing, cells were harvested for bulk mRNA-sequencing and ICP0 immunofluorescence staining. b, Relative densities of the percentage of viral transcripts (log10 transformed) per cell for the three time points post infection. c, Relative densities of the percentage of viral transcripts per cell (log10 transformed) for G1 and non-G1 cells for cells harvested at 3hpi and 5hpi. d, Heatmap of expression values of the first eight expressed viral genes for experiment two with only “HSV-1 high” cells. Rows (genes) and columns (cells) were sorted as described in the main text for Figure 2. Above the heatmap, cell cycle and log10 transformed percentage of viral transcripts are shown. e, Relationship between differentially expressed genes and viral transcription. The horizontal axis shows the maximal log(2) transformed bulk RNA-seq fold change of the three time points after infection compared to uninfected cells, the vertical axis the linear correlation coefficient of the gene expression with the sum of viral transcripts in bins of 20 cells, using only “HSV-1 high” cells. f, Cells were projected on a two-dimensional map using UMAP according to gene expression values, with cells colored by harvesting time point. Main distinguishing features were marked in grey. g, Coloring by amount of HSV-1 transcripts. Cells without HSV-1 transcripts are colored in light gray. h, cells colored by expression values (top panels) and RNA velocity values (bottom panels) of the indicated genes. Color scales are shown to the right of these panels. For expression values, areas containing cells with relatively high expression levels were marked with a light green ellipse. The dark green and blue ellipse denote S and G2/M phase cells, respectively. For RNA velocity, cells for which no value could be calculated were colored in light gray. i, Correlation of gene expression with NQO1 expression (horizontal axis) and SULF1 expression (vertical axis) in cells harvested at 3hpi. Every dot represents a gene, colored by the slope of the linear correlation with NQO1. Selected genes were labeled by name. Note that gene names were slightly jittered to improve readability. j, Groups of cells according to PAGA were labeled A to S. Transition probabilities between clusters are proportional to the line thickness between the clusters. Areas of interest are marked with light green ellipses.

Figure S6 | a, HEK 293 cells were treated for 16h with solvent (DMSO) or different concentrations of Bardoxolone methyl, followed by RNA isolation. NQO1 mRNA levels were probed using RT-qPCR. b, HEK 293 cells were infected with HSV-1 at an MOI of 1. After removal of virus inoculum, washing with PBS, and adding back conditioned mediums, solvent or different concentrations of Bardoxolone methyl were added. At 16hpi, viral DNA was probed in the supernatant using qPCR (left panel) and viral mRNAs in the RNA isolated from the cells using RT-qPCR (right panel). c, Levels of NQO1 mRNA in infected cells from Fig. 6b and S6b. d, levels of TOP2A and GMNN mRNAs in cells treated with Bardoxolone methyl from Fig. 6a and S6a. For all panels, barplots indicate means, error bars denote standard deviations, the individual measurement values are shown as grey dots.

## Material and Methods

### Cells and virus

Primary normal human dermal fibroblasts (PromoCell C-12300) were cultured in Dulbecco’s modified Eagle medium (DMEM) supplemented with 10% fetal bovine serum (FBS), 100 units/ml penicillin, 100 μg/ml streptomycin and 1%NAEE at 37°C and 5% CO2. Primary cells were cultured until Passage 10. Wild-type HSV-1 was derived from a patient isolate and was obtained from Department of Virology, Saarland University Medical School, Homburg, Germany. The virus was propagated in NHDF cells. At 72 hpi, viral supernatant was collected and sterile-filtered through a 0.45 μm pore size filter and stored at −80°C. Viral titers were determined by plaque assay as described previously [5].

### Infections

Cells were seeded in 10 cm dishes (for scRNA-seq) and in 6 well plates (for bulk RNA samples and slides for immunofluorescence). For the non-synchronized infection at 37 °C, half of the medium (conditioned medium) was removed, and then cells were incubated for 30 min with HSV-1 (MOI 10) or were mock infected. The supernatant was removed and cells were washed with PBS, followed by re-applying the conditioned medium. We harvested uninfected cells and at early time points (1hpi, 3hpi, and 5hpi) in two biological replicates.

Following the same protocol of infection for synchronized infection, cells were incubated for 20 min at 4°C prior to infection. Virus containing supernatant was incubated for 1 h also at 4°C for 1h. After inoculation cells were washed with PBS and then complete Fresh DMEM Media was applied and cultured at 37°C. For Drop-Seq all cells were washed with cold PBS and fixed in ice cold Methanol (80%).

### DropSeq single-cell RNA-sequencing

Cells were fixed and used for Drop-seq single-cell sequencing as previously described [24]. Monodisperse droplets of about 1 nl in size were prepared on a self-built Drop-seq set up following closely the instrument set up and library generation procedure as described [23]. Principle: Upon nanoliter droplet formation, individual cells are co-encapsulated with individual, uniquely barcoded beads, and become lysed. Released polyadenylated RNA molecules then hybridize to polyd(T) primers that are attached to the uniquely barcoded beads. Each captured mRNA molecule is tagged with a barcode indicating its cell of origin and a unique molecular identifier. Nanoliter-droplets are collected and broken, and RNA molecules are reverse transcribed into complementary DNA (cDNA), amplified by PCR, size-selected for mRNA, fragmented and 3’ ends sequenced in bulk.

cDNA libraries had large average sizes of about 1.5 kb, indicating high quality RNA and cDNA molecules. 600 pg of each cDNA library was fragmented and amplified for sequencing with the Nextera XT v2 DNA Library Preparation kit (Illumina, San Diego, CA, USA). Single-cell libraries at 1.8 pM (final average insert sizes ranging from 580 to 680 bp) were sequenced in paired-end mode on Illumina Nextseq 500 sequencers using Illumina Nextseq500/550 High Output v2 kits (75 cycles). Read 1: 20 bp (bases 1-12 cell barcode, bases 13-20 UMI; Dropseq custom primer 1 “Read1CustSeqB”), index read: 8 bp, read 2 (paired end): 64 bp).

### Bulk RNA-sequencing and analysis

RNA was extracted from cells using Trizol (Thermo Fisher) and the RNA clean and concentrator kit (Zymo). Sequencing libraries were prepared using the TruSeq Stranded mRNA kit (Illumina) according to the manufacturer’s instruction and sequenced on a HiSeq4000 device using 2×76nt paired-end sequencing. Sequencing reads were aligned to the hg38 version of the human genome using tophat2 [64]. Standard differential expression analysis was performed using quasR [65] for counting reads (using the Gencode gene annotation version 28) and edgeR [66].

### Single-cell RNA-seq data processing

The scRNA-seq data was processed using the PiGx pipeline [67]. In short, polyA sequences are removed from reads. The reads are mapped to the hg38 version of the human genome using STAR [68] with gene models from Gencode version 28. The number of cells, for each sample, is determined using dropbead [24]. Finally, a combined digital expression matrix is constructed, containing all sequenced experiments. This table is available at the GEO entry. Cells containing less than 2000 detected genes were filtered from the analysis. Normalized counts and tSNE coordinates were calculated using the Seurat package [69]. Cell cycle was assigned to each cell using the CellCycleScoring function. Cell cycle gene sets were taken from [70]. Normalized total HSV-1 transcription was calculated by summing up raw counts of all detected viral genes, divided by the total number of raw counts, multiplied by a scaling factor of 10000 and log(2) transformed. Plots were generated using ggplot2 [71], pheatmap or ComplexHeatmap [72].

### Diffusion maps for Herpes

Transition probabilities between cells were estimated based on the diffusion pseudo-time distances [29], implemented in the destiny Bioconductor package [28]. Diffusion pseudo-time distance between two cells represents the cumulative probability of travelling from one cell to the other on a probabilistic graph. The graph is constructed by embedding cells using gaussian kernels. Obtained distances were converted to transition probabilities using an exponential probability distribution.

### Binning and correlations

Correlations of host cell gene expression with HSV-1 gene expression (Fig. 4), SULF1 or NQO1 expression (Fig. 5) were calculated using binned cells to reduce noise. For this, cells with “high” expression (see Fig. S3a and S4c) were first sorted according to normalized values of HSV-1 transcripts, SULF1 or NQO1. Then, cells were binned with 20 cells per bin. Each bin was then treated as a “metacell”. Normalized gene expression values for the metacella were calculated by summing up, per gene, all raw values in the bin, dividing by the total raw count in the entire metacell, multiplied by a scaling factor of 10000 and log(2) transformed. Then, the linear Pearson correlation coefficient and slope for each gene or antisense transcript group with HSV-1 transcripts, SULF1 or NQO1, respectively, were calculated.

### RNA interference

NHDF cells were plated in 6-well plates and incubated at 37°C. 24 hours later cells were transfected with siRNAs (final amount 15 pmol) using Lipofectamine® RNAiMax (Thermo Fischer) according to the manufacturer’s instructions. Briefly, siRNAs was diluted in 125μL of reduced serum medium (OPTI-MEM I; Invitrogen). The Lipofectamine RNAiMax reagent was diluted in 125μL of OPTI-MEM I and the two solutions were then mixed and incubated for 10 minutes at room temperature before addition to the cells. 8 hours after transfection, cells were infected with HSV 1 (MOI 1) as mentioned above. After that cells were again transfected with the siRNAs and RNA Samples and cell supernatants were harvested 16 hours after infection. SiRNAs that were used in this study to transfect NHDF cells are described in supplementary table S4.

### Measuring viral DNA in cell culture supernatant

Cell culture supernatant was treated with 2 mg/ml proteinase K (Thermo Fisher) in 30 mM Tris pH 8 and 0.5% Triton for 10 minutes at 70 °C, followed by inactivation of the proteinase for 10 minutes at 95 °C. After 1:1 dilution in water, the samples were used for qPCR using the UL29 primer pair (supplementary table S4). A standard curve for relative quantification was prepared from serial dilution of a virus stock with 1 PFU/ml (measured using plaque assays) treated in parallel to the samples. For all experiments, four measurements were done for two biological replicates.

### RT-qPCR

RNA was extracted from cells using Trizol (Thermo Fisher) and the RNA clean and concentrator kit (Zymo). DNase treatment using DNase I amplification grade (Thermo Fisher) and RT using SuperScript III (ThermoFisher) was performed according to the manufacturer’s protocol. For all experiments, four measurements were done from two biological replicates. Power SYBR Green PCR Master Mix on a StepOnePlus system (both Thermo Fisher Scientific) was used for qPCR. Data were normalized to the indicated timepoint/sample and the GAPDH signal using the ΔΔCt method [73]. Primers for qPCR were designed using the Universal ProbeLibrary (Roche Life Sciences), tested for efficiency and a single amplicon and listed in supplementary table S4.

### UMAP

For visualization purposes, we employed in Fig. 4 and 5 the UMAP dimensionality reduction algorithm [74] instead of tSNE as in the previous figures. The reason was that the directionality, which the RNA velocity algorithm calculates based on all detectable genes, and then projects as arrow clouds on the two-dimensional map (Fig. 4C), could not be visualized when using tSNE. Variable genes were define using the FindVariableGenes function with the default parameters. First twenty principle components were calculated using the defined variable genes. Dimensionality reduction was performed with UMAP [74]. The cells were clustered using Louvain algorithm with a set of resolution parameters ranging from 0.5 – 2.

### Velocity and graph abstraction

The data was pre-processed using the Velocyto CLI pipeline. Only cells with more than 500 intronic and more than 500 exonic reads were kept for further analysis. Intronic signal represented about 3% of all sequencing reads.

The following parameters were used within the Velocyto analysis:

vlm.score_detection_levels()

vlm.filter_genes(by_detection_levels=True)

vlm.score_cv_vs_mean(2000, max_expr_avg=55)

vlm.filter_genes(by_cv_vs_mean=True)

vlm.normalize_by_total()

vlm.pca = pca(n_components=10, svd_solver=’arpack’, random_state=1)

vlm.pcs = vlm.pca.fit_transform(vlm.S_norm.T)

vlm.knn_imputation(k=100, balanced=True, b_sight=3000, b_maxl=1500, n_jobs=16)

vlm.fit_gammas(weighted=False)

vlm.estimate_transition_prob(hidim=“Sx_sz”, embed=”ts”, transform=’log’)

vlm.calculate_embedding_shift(sigma_corr = 0.05, expression_scaling=False)

vlm.calculate_grid_arrows(smooth=0.8, steps=(40, 40), n_neighbors=100)

For the nascent RNA clustering, the Velocyto estimated transcriptional rates were processed using scanpy [75]. The rates were used to calculate the principal components, UMAP embeddings, cluster the cells using the Louvain algorithm, and compute PAGA [38].

### Bardoxolone methyl treatment

Bardoxolone methyl was purchased from MedChemExpress (HY-13324). Stock concentrations of 100 and 400 μM were prepared in DMSO, and directly added to the cell culture medium. For the assays combining Bardoxolone methyl treatment with HSV-1 infection, half of the medium was removed, virus inoculum at an MOI of 1 was added to the remaining medium, and half an hour later the cells were washed with warm PBS, the conditioned medium added back to the cells, and then Bardoxolone methyl at the indicated concentrations supplied.

