## Supplementary Material for "Single-cell RNA-sequencing of Herpes simplex virus 1-infected cells identifies NRF2 activation as an antiviral program"

Figure S1

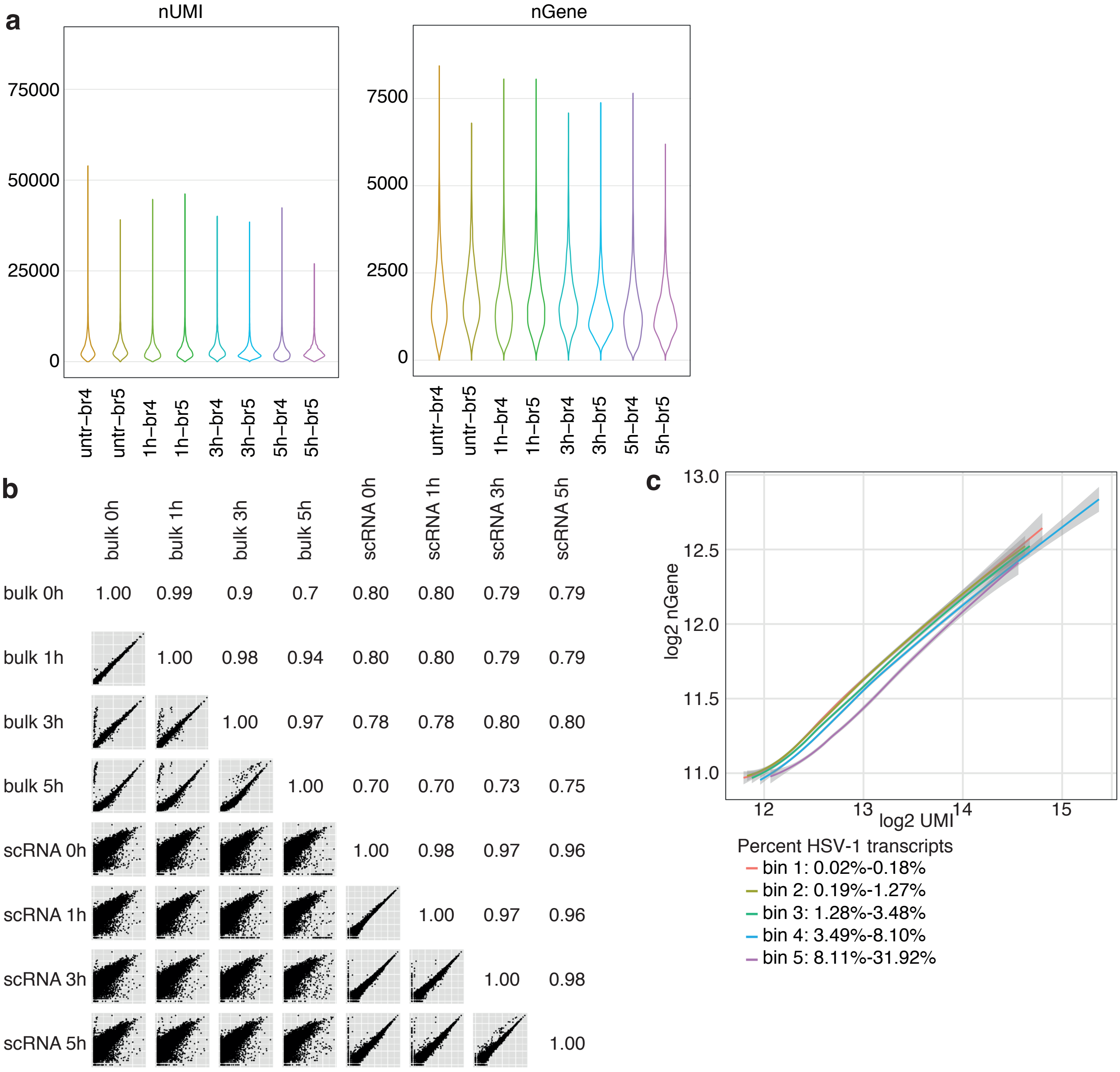

**Figure S1** | a, Violin plots showing the distribution of number of unique molecular identifiers and number of detected genes in the individual scRNA-seq samples. b, Correlation of bulk RNA-seq with scRNA-seq data. Shown are scatter plots (lower left part) and linear correlation coefficients (upper right part) of bulk polyA RNA-seq values (horizontal axes, log(2) transformed fpkm values) and scRNA-seq values (vertical axes, log(2) transformed sums of scRNA-seq counts) of merged replicates, c, Cells harvested at 5hpi containing at least one viral transcript were binned according to the percentage of viral transcripts per cell, and the log2 transformed nUMI/nGene values plotted as smoothed means with 95% confidence intervals in gray. Percentage ranges of the bins are indicated below the plot.

### Figure S2, part1

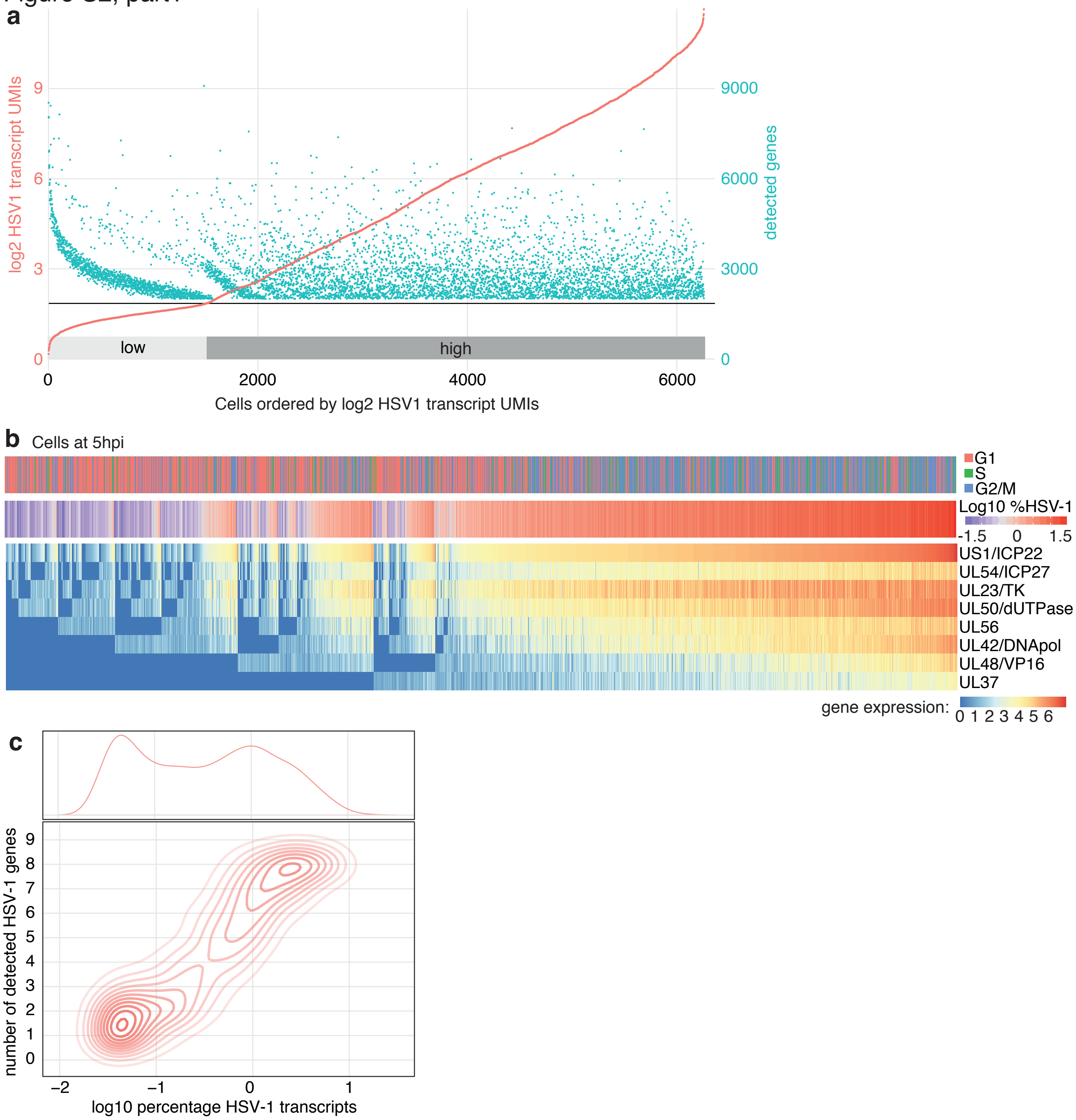

**Figure S2** I a, Relationship between normalized levels of HSV-1 transcripts and number of detected genes shows two categories of cells. Cells with at least one viral transcript were sorted by normalized HSV-1 transcript UMIs. For each cell, the red dot represents the log(2) transformed sum of HSV-1 transcript UMIs (left axis), and the green dot the number of genes detected (right axis). Note that only cells with more than 2000 detected genes were used, thus the lower apparent limit for the green dots. Light and dark gray bars designate "low HSV-1" cells and "high HSV-1" cells according to the bimodality of the plot. The horizontal gray line denotes the cutoff expression between high and low. For characterization of the gene expression cascade of viral transcripts, only the 3896 "high HSV-1" cells were used, in order to reduce the sampling error caused by the detection rate. b, Heatmap of expression values of the first eight expressed viral genes for experiment two with only "HSV-1 high" 5hpi cells as defined in the main text. Rows (genes) and columns (cells) were sorted as described in the main text for Figure 3b. Above the heatmap, cell cycle and log10 transformed percentage of viral transcripts are shown. c, Smoothened two-dimensional densities of the number of detected viral genes (vertical axis) in relationship to the percentage of viral transcripts (horizontal axis) for the set of cells used for Fig. 2b. Relative densities as in Fig. 2b are shown on top. d, Clustered heatmap of transition probabilities on the diffusion maps of cells bearing HSV-1 transcripts. Left part, clustered probabilities colored from white (low) to red (high). Columns and rows represent cells in the same order. Right part, expression of the first eight viral genes. The dotted rounded rectangle represents the cluster of transition probabilities between the states denoted under the heatmap. For this analysis, only S/G2/M phase cells were used, since the difference between G1 and non-G1 single-cell transcriptomes otherwise dominates the transition probabilites. Also, cells harvested at 5hpi are left out in order to better study early viral gene expression.

Figure S2, part 2

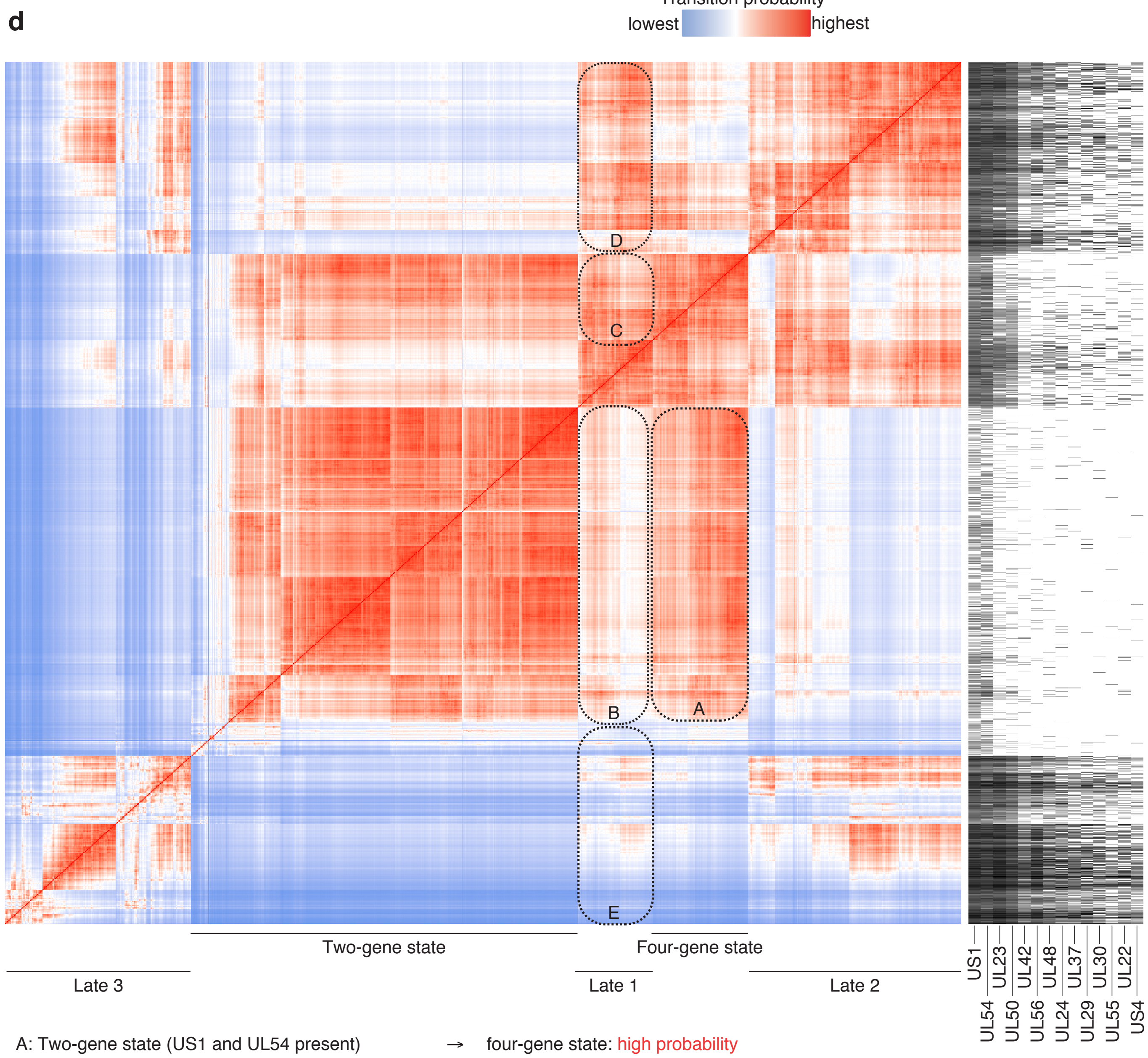

A: Two-gene state (US1 and UL54 present) → four-gene state: high probability
B: Two-gene state (US1 and UL54 present) → late 1: low probability
C: Four-gene state (US1/UL54/UL23/UL50 present) → late 1: high probability
D: Late 1 → late 2: high probability
E: Late 1 → late 3: low probability

### Figure S3

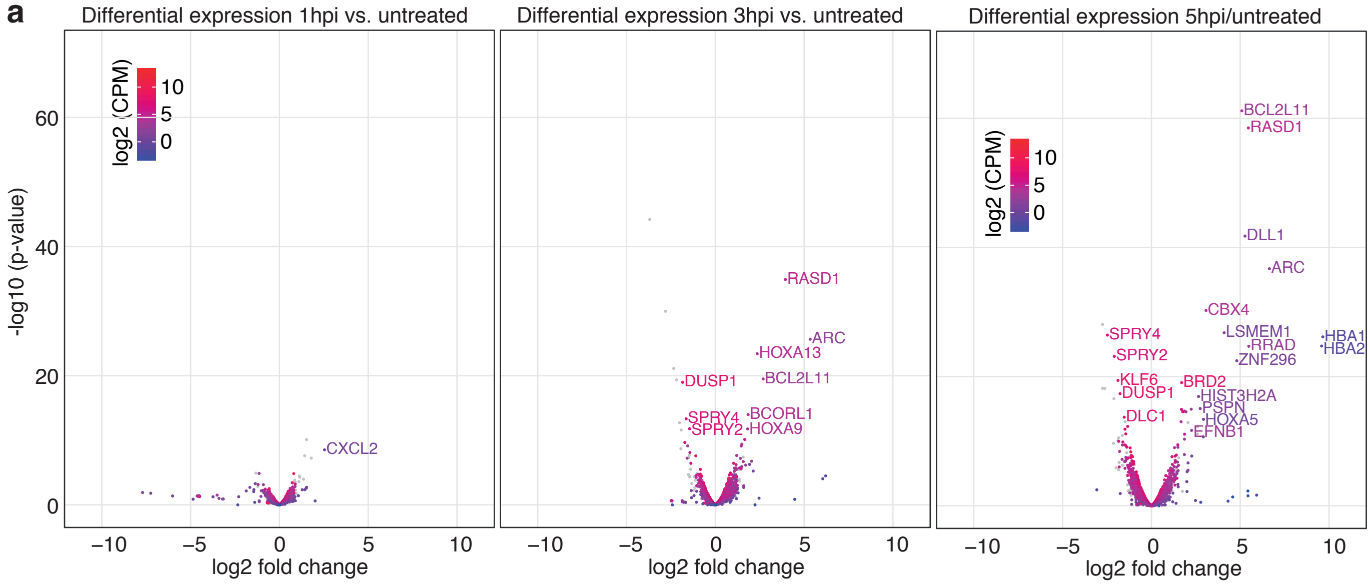

**Figure S3** I a, Differential bulk mRNA expression comparing expression from 1hpi (left panel), 3hpi (middle panel), and 5hpi (right panel) with untreated cells. Outliers were marked by name. Horizontal axis denote log(2) transformed fold-changes, vertical axis log(10) transformed p-values. The color represents the log(2) transformed counts per millions over all time points. Low-reproducibility genes (see Supplementary Table S3) are displayed as light gray dots.

#### Figure S4

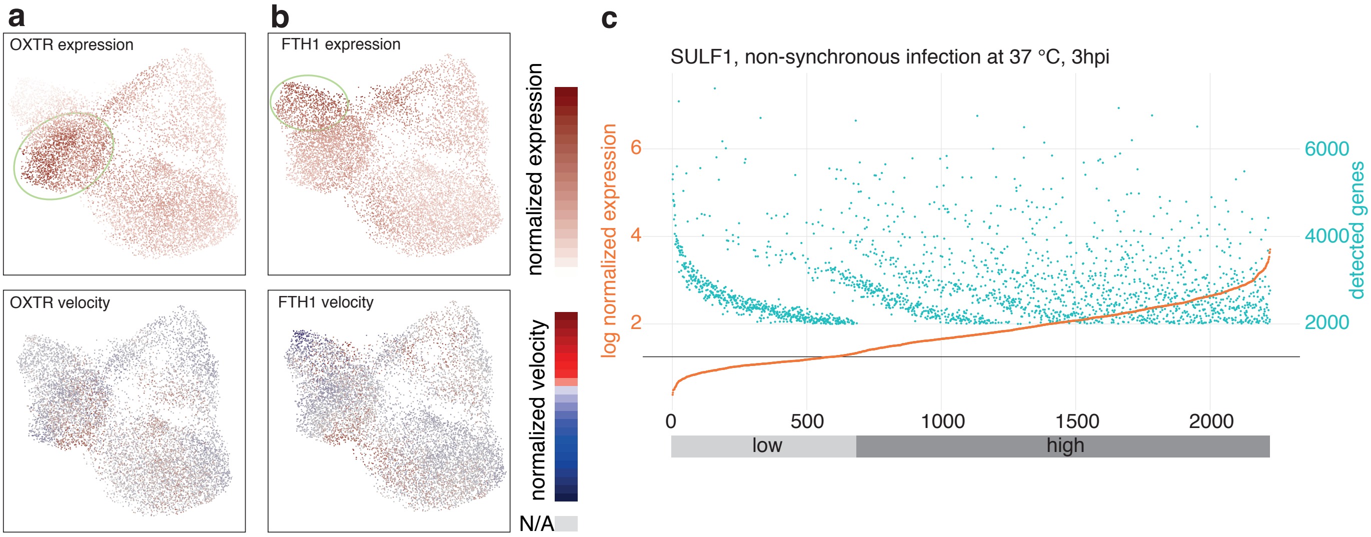

**Figure S4** I ab, cells colored by expression values (top panels) and RNA velocity values (bottom panels) of the indicated genes. Color scales are shown to the right of these panels. For expression values, areas containing cells with relatively high expression levels were marked with a light green ellipse. For RNA velocity, cells for which no value could be calculated were colored in light gray. c, Cells harvested at 3hpi were sorted by normalized SULF1 transcript UMIs. For each cell, the red dot represents the log(2) transformed amount of SULF1 transcript UMIs (left axis), and the green dot the number of genes detected (right axis). Note that only cells with more than 2000 detected genes were used, thus the lower apparent limit for the green dots. Light and dark gray bars designate low SULF1 cells and high SULF1 cells according to the bimodality of the plot. The horizontal gray line denotes the cutoff expression between high and low. Only high SULF1 cells were used to calculate the correlations in Fig. 4h.

Figure S5, part1

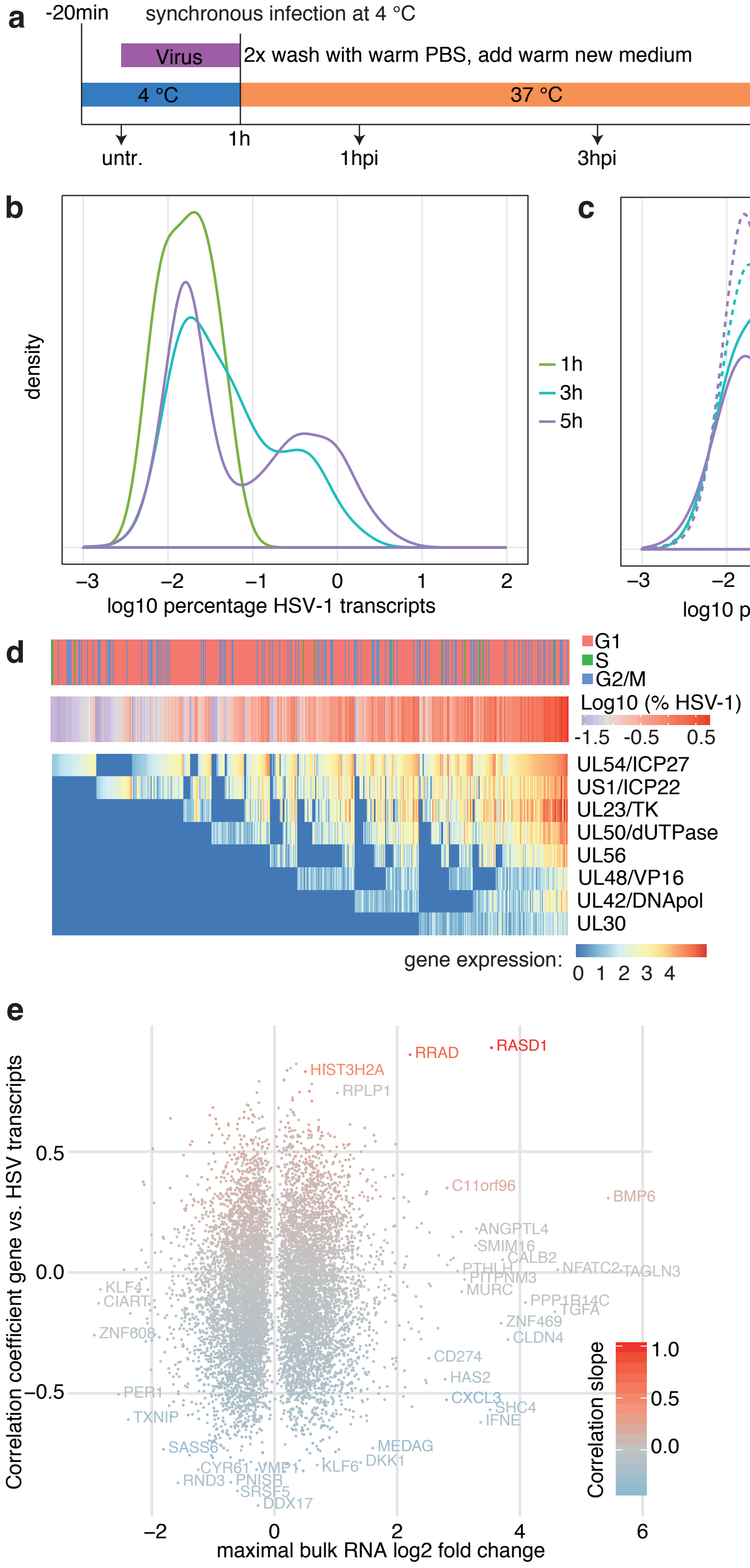

**Figure S5** I a, Infection protocol for the synchronized infection at 4 °C. In parallel to single-cell RNA-sequencing, cells were harvested for bulk mRNA-sequencing and ICP0 immunofluorescence staining. b, Relative densities of the percentage of viral transcripts (log10 transformed) per cell for the three time points post infection. c, Relative densities of the percentage of viral transcripts per cell (log10 transformed) for G1 and non-G1 cells for cells harvested at 3hpi and 5hpi. d, Heatmap of expression values of the first eight expressed viral genes for experiment two with only "HSV-1 high" cells. Rows (genes) and columns (cells) were sorted as described in the main text for Figure 2. Above the heatmap, cell cycle and log10 transformed percentage of viral transcripts are shown. e, Relationship between differentially expressed genes and viral transcription. The horizontal axis shows the maximal log(2) transformed bulk RNA-seq fold change of the three time points after infection compared to uninfected cells, the vertical axis the linear correlation coefficient of the gene expression with the sum of viral transcripts in bins of 20 cells, using only "HSV-1 high" cells. f, Cells were projected on a two-dimensional map using UMAP according to gene expression values, with cells colored by harvesting time point. Main distinguishing features were marked in grey. g, Coloring by amount of HSV-1 transcripts. Cells without HSV-1 transcripts are colored in light gray. h, cells colored by expression values (top panels) and RNA velocity values (bottom panels) of the indicated genes. Color scales are shown to the right of these panels. For expression values, areas containing cells with relatively high expression levels were marked with a light green ellipse. The dark green and blue ellipse denote S and G2/M phase cells, respectively. For RNA velocity, cells for which no value could be calculated were colored in light gray. i, Correlation of gene expression with NQO1 expression (horizontal axis) and SULF1 expression (vertical axis) in cells harvested at 3hpi. Every dot represents a gene, colored by the slope of the linear correlation with NQO1. Selected genes were labeled by name. Note that gene names were slightly jittered to improve readability. j, Groups of cells according to PAGA were labeled A to S. Transition probabilities between clusters are proportional to the line thickness between the clusters. Areas of interest are marked with light green ellipses.

Figure S5, part 2

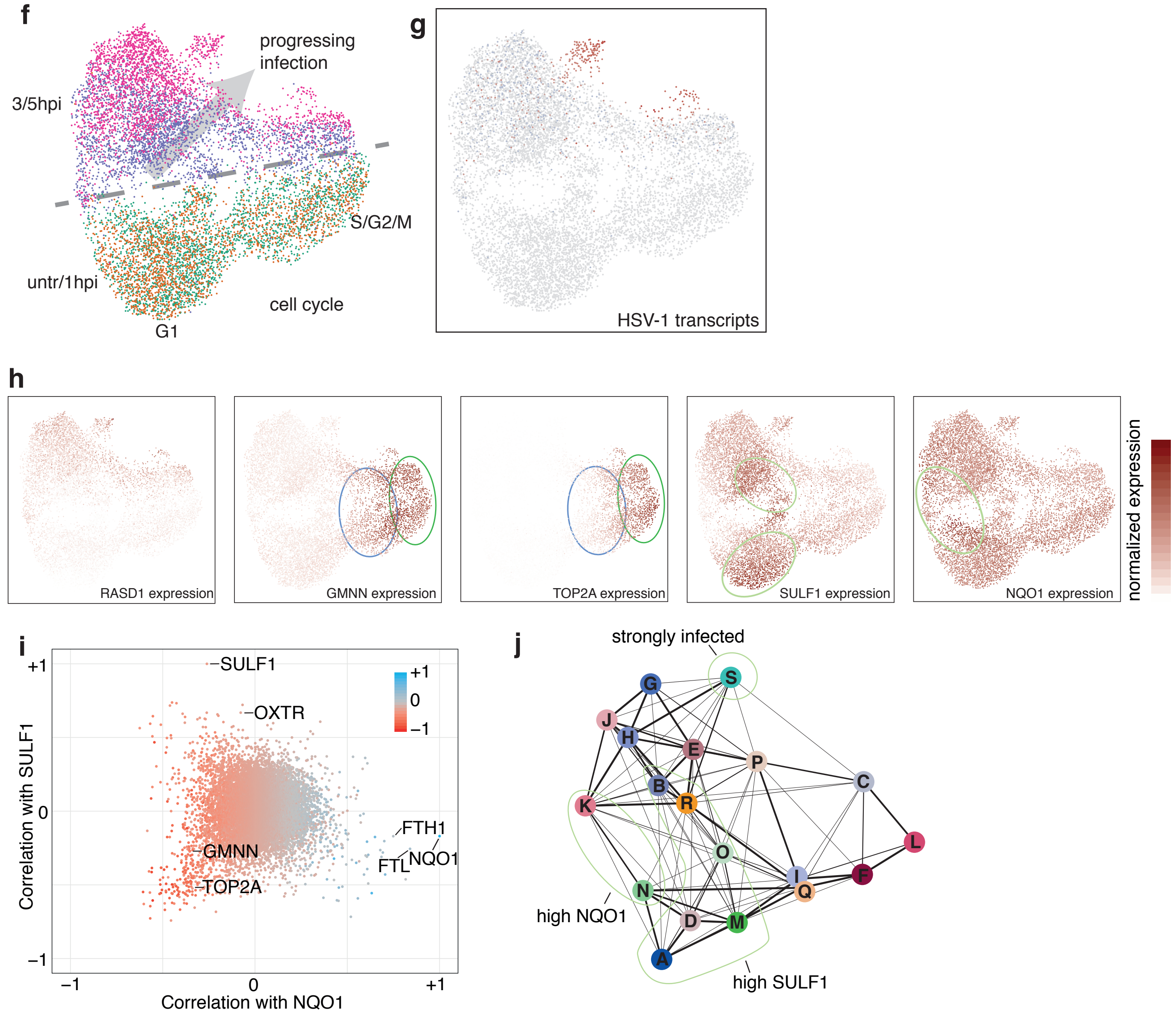

### Figure S6

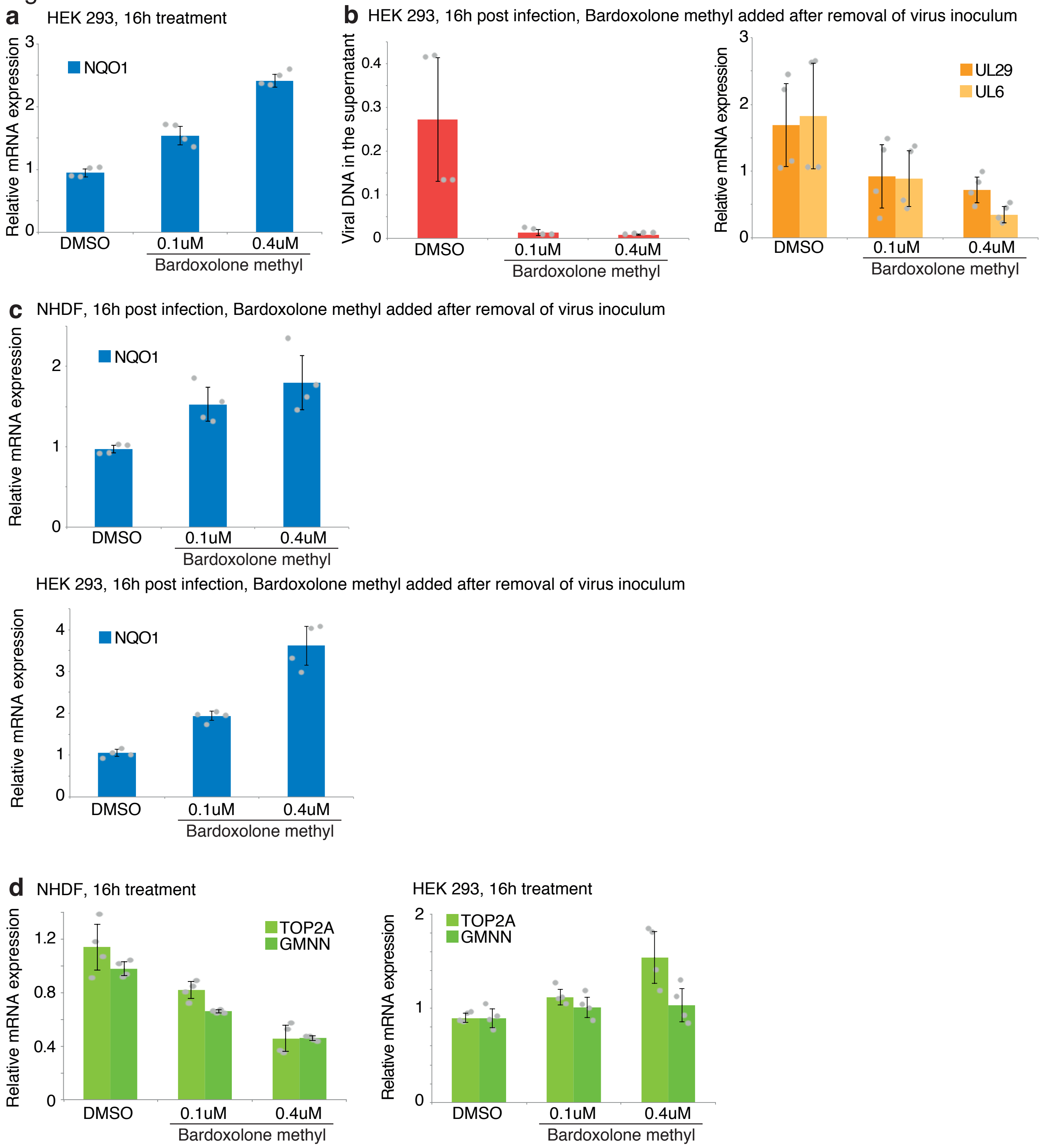

**Figure S6** I a, HEK 293 cells were treated for 16h with solvent (DMSO) or different concentrations of Bardoxolone methyl, followed by RNA isolation. NQO1 mRNA levels were probed using RT-qPCR. b, HEK 293 cells were infected with HSV-1 at an MOI of 1. After removal of virus inoculum, washing with PBS, and adding back conditioned mediums, solvent or different concentrations of Bardoxolone methyl were added. At 16hpi, viral DNA was probed in the supernatant using qPCR (left panel) and viral mRNAs in the RNA isolated from the cells using RT-qPCR (right panel). c, Levels of NQO1 mRNA in infected cells from Fig. 6b and S6b. d, levels of TOP2A and GMNN mRNAs in cells treated with Bardoxolone methyl from Fig. 6a and S6a. For all panels, barplots indicate means, error bars denote standard deviations, the individual measurement values are shown as grey dots.

**Table S1: Samples**

**Pilot experiment (not analyzed in manuscript): Replicate 1, synchronized 4 °C infection**

| | |
|---|---|
| *Replicate 1 - 0h (untreated cells)* | scRNA-seq Rep 1 0hpi |
| *Replicate 1 - 3hpi* | scRNA-seq Rep 1 3hpi |
| *Replicate 1 - 5hpi* | scRNA-seq Rep 1 5hpi |

**Replicate 2 and 3, synchronized 4 °C infection**

| | | | |
|---|---|---|---|
| *Replicate 2 - 0h (untreated cells)* | scRNA-seq Rep 2 0h | bulk polyA RNA-seq Rep 2 0h | ICP0 Immunofluorescence (not shown) |
| *Replicate 2 - 1hpi* | scRNA-seq Rep 2 1h | bulk polyA RNA-seq Rep 2 1h | ICP0 Immunofluorescence (not shown) |
| *Replicate 2 - 3hpi* | scRNA-seq Rep 2 3h | bulk polyA RNA-seq Rep 2 3h | ICP0 Immunofluorescence (not shown) |
| *Replicate 2 - 5hpi* | scRNA-seq Rep 2 5h | bulk polyA RNA-seq Rep 2 5h | ICP0 Immunofluorescence (Fig. S2B) |
| *Replicate 3 - 0h (untreated cells)* | scRNA-seq Rep 3 0h | bulk polyA RNA-seq Rep 3 0h | ICP0 Immunofluorescence (not shown) |
| *Replicate 3 - 1hpi* | scRNA-seq Rep 3 1h | bulk polyA RNA-seq Rep 3 1h | ICP0 Immunofluorescence (not shown) |
| *Replicate 3 - 3hpi* | scRNA-seq Rep 3 3h | bulk polyA RNA-seq Rep 3 3h | ICP0 Immunofluorescence (not shown) |
| *Replicate 3 - 5hpi* | scRNA-seq Rep 3 5h | bulk polyA RNA-seq Rep 3 5h | ICP0 Immunofluorescence (not shown) |

**Replicate 4 and 5, non-synchronized 37 °C infection, conditioned medium**

| | | | |
|---|---|---|---|
| *Replicate 4 - 0h (untreated cells)* | scRNA-seq Rep 4 0h | bulk polyA RNA-seq Rep 4 0h | ICP0 Immunofluorescence (not shown) |
| *Replicate 4 - 1h* | scRNA-seq Rep 4 1h | bulk polyA RNA-seq Rep 4 1h | ICP0 Immunofluorescence (not shown) |
| *Replicate 4 - 3h* | scRNA-seq Rep 4 3h | bulk polyA RNA-seq Rep 4 3h | ICP0 Immunofluorescence (not shown) |
| *Replicate 4 - 5h* | scRNA-seq Rep 4 5h | bulk polyA RNA-seq Rep 4 5h | ICP0 Immunofluorescence (Fig. S1B) |
| *Replicate 5 - 0h (untreated cells)* | scRNA-seq Rep 5 0h | bulk polyA RNA-seq Rep 5 0h | ICP0 Immunofluorescence (not shown) |
| *Replicate 5 - 1h* | scRNA-seq Rep 5 1h | bulk polyA RNA-seq Rep 5 1h | ICP0 Immunofluorescence (not shown) |
| *Replicate 5 - 3h* | scRNA-seq Rep 5 3h | bulk polyA RNA-seq Rep 5 3h | ICP0 Immunofluorescence (not shown) |
| *Replicate 5 - 5h* | scRNA-seq Rep 5 5h | bulk polyA RNA-seq Rep 5 5h | ICP0 Immunofluorescence (not shown) |
| *Mock - 0h (untreated cells)* | | bulk polyA RNA-seq Mock 0h | |
| *Mock - 1h* | | bulk polyA RNA-seq Mock 1h | |
| *Mock - 3h* | | bulk polyA RNA-seq Mock 3h | |
| *Mock - 5h* | | bulk polyA RNA-seq Mock 5h | |

**Table S2: quantified cells with more than 500/2000 detected genes**

**Pilot experiment: Replicate 1, synchronized 4 °C infection**

| | **0 hpi** | **1 hpi** | **3 hpi** | **5 hpi** | **Sum per row** |
|---|---|---|---|---|---|
| Replicate 1 | 3713 | N/A | 3634 | 8370 | 15717 |

**quantified cells with more than 2000 genes detected**

| | **0 hpi** | **1 hpi** | **3 hpi** | **5 hpi** | **Sum per row** |
|---|---|---|---|---|---|
| Replicate 1 | 888 | N/A | 1308 | 1508 | 3704 |
| With HSV-1 genes | 0 | N/A | 72 | 482 | |
| Percent with HSV-1 genes | 0 | 0 | 5,50% | 31,96% | |

**Replicate 2 and 3, synchronized 4 °C infection**

| | **0 hpi** | **1 hpi** | **3 hpi** | **5 hpi** | **Sum per row** |
|---|---|---|---|---|---|
| Replicate 2 | 4409 | 5106 | 4500 | 4850 | 18865 |
| Replicate 3 | 5802 | 6681 | 5640 | 5766 | 23889 |
| Sum per column | 13924 | 11787 | 13774 | 18986 | 58471 |
| With HSV-1 genes | 0 | 0 | 440 | 4423 | |

**quantified cells with more than 2000 genes detected**

| | **0 hpi** | **1 hpi** | **3 hpi** | **5 hpi** | **Sum per row** |
|---|---|---|---|---|---|
| Replicate 2 | 1158 | 1300 | 1520 | 1405 | 5234 |
| Replicate 3 | 2131 | 1642 | 1795 | 1495 | 6960 |
| Sum per column | 3289 | 2942 | 3315 | 2900 | 15764 |
| With HSV-1 genes | 0 | 5 | 134 | 870 | |
| Percent with HSV-1 genes | 0 | 0,17% | 4,04% | 30,00% | |

**Replicate 4 and 5, non-synchronized 37 °C infection, conditioned medium**

| | **0 hpi** | **1 hpi** | **3 hpi** | **5 hpi** | **Sum per row** |
|---|---|---|---|---|---|
| Replicate 4 | 4409 | 5106 | 4500 | 4850 | 18865 |
| Replicate 5 | 5802 | 6681 | 5640 | 5766 | 23889 |
| Sum per column | 13924 | 11787 | 13774 | 18986 | 58471 |
| With HSV-1 genes | 0 | 0 | 440 | 4423 | |

**quantified cells with more than 2000 genes detected**

| | **0 hpi** | **1 hpi** | **3 hpi** | **5 hpi** | **Sum per row** |
|---|---|---|---|---|---|
| Replicate 4 | 1813 | 1858 | 1601 | 1030 | 6302 |
| Replicate 5 | 1993 | 1887 | 1166 | 973 | 6019 |
| Sum per column | 3806 | 3745 | 2767 | 2003 | 12321 |
| With HSV-1 genes | 0 | 450 | 2238 | 2001 | |
| Percent with HSV-1 genes | 0 | 12,02% | 80,88% | 99,90% | |

**Table 3: Omitted genes**

| Mock variable genes | Variable between experiments |
|---|---|
| ANKRD9 | ABHD17A |
| ARL4D | ACAN |
| ARL5B | ACP5 |
| ATRX | ACTC1 |
| BACH1 | ADAMTS4 |
| BBX | ADAT3 |
| BTG2 | ADCK5 |
| C2orf44 | AGAP2 |
| CDK6 | AGRN |
| CENPA | AKR1B1 |
| CLOCK | AKR1B10 |
| CSRNP1 | ALDH16A1 |
| CXCL1 | ALDH1A1 |
| CXCL8 | ALDH1A3 |
| DDIT4 | AMDHD2 |
| DST | ANKRD1 |
| DUSP6 | ANKRD33B |
| EEA1 | AP5Z1 |
| EGR1 | APBA2 |
| ERRFI1 | APOE |
| FAM111B | AREG |
| FAM63B | ARHGAP28 |
| FCHO2 | ASPHD1 |
| FILIP1L | ATAD3A |
| GCC2 | ATAD3B |
| HBEGF | ATF3 |
| HES1 | BAALC |
| HIST2H3D | BHLHE40 |
| IER3 | BMP4 |
| IL11 | BOLA2 |
| INHBA | BOLA2B |
| JMJD1C | BOP1 |
| JUNB | BORCS6 |
| KIAA1109 | BRAT1 |
| KRTAP2-3 | C10orf10 |
| LINC00506 | C10orf107 |
| LOC102724238 | C10orf55 |
| LOC554249 | C14orf80 |
| LOC728673 | C16orf59 |
| LPP | C19orf24 |
| MOB1B | C19orf71 |
| NEK2 | C1orf159 |
| NEURL1B | C4orf48 |
| NFAT5 | CAPN15 |
| NIPBL | CARD9 |
| NKTR | CASS4 |
| NR4A1 | CCDC144B |
| NRIP1 | CD200 |
| PIK3C2A | CD82 |
| PLCB4 | CDCP1 |
| POLK | CDKN3 |
| PRKG1 | CDON |
| PTGS2 | CDT1 |
| RANBP2 | CHTF18 |
| RGPD2 | CLUH |
| RGPD3 | CLUL1 |
| RGPD4 | COL13A1 |
| RGPD8 | COL18A1 |
| ROCK2 | COL4A1 |
| RPLP0P2 | COL5A3 |
| RPS14P3 | COL6A1 |
| SACS | COL6A2 |
| SIAE | COL7A1 |
| SLC7A11 | CORO7 |
| SMC5 | CPA4 |
| SREK1 | CSPG4 |
| STAG3L1 | CTB-113P19.1 |
| STAG3L3 | CTC-436P18.1 |
| TAOK1 | CTSD |
| THOC2 | CTU2 |
| TMF1 | CXCL14 |
| UFL1 | CXCL5 |
| USP34 | CXCL6 |
| USP53 | CYBA |
| USP9Y | CYP1B1 |
| UTRN | DES |
| VGLL3 | DGKQ |
| YOD1 | DHRS3 |
| ZC3H13 | DIRAS3 |
| ZEB1 | DLL4 |
| ZSCAN26 | DOHH |
| | DOK5 |
| | DUS3L |
| | DUSP2 |
| | DUSP5 |
| | DVL1 |

E2F1
EDIL3
EDN1
EFCAB7
EGFL7
EGR1
EGR2
EGR3
EMILIN1
EPB41L3
EPHA4
EPHB1
EPHB2
EVA1B
F2RL1
FAM173A
FAM196B
FAM65B
FAM71E1
FASN
FBXL15
FBXL6
FHL1
FIBCD1
FILIP1L
FLNC
FMO4
FOS
FOSB
G0S2
GABBR2
GADD45B
GAL
GALNT18
GCH1
GCNT4
GDF9
GMPPB
GNB1L
GPC1
GPR1
GPR68
GTPBP6
HAGHL
HCG4B
HES1
HES4
HIST1H2AC
HIST1H2BJ
HIST1H4E
HIST2H2AA3
HIST2H2AA4
HIST2H2AC
HIST2H3D
HMGN2P46
HOXD4
HR
HS3ST3A1
HSPA1A
HSPA1B
IFIT1
IFIT2
IGFBP2
IL24
IL33
INF2
IRAK1BP1
ISG15
ITGA2
JOSD2
KCNC4
KCNJ3
KCTD12
KISS1
KLF10
KLF9
KLHL17
KRBOX1
KRTAP2-3
LACAT8
LCE2A
LIF
LIN7A
LINC00957
LINC00968
LINC01133
LINC01336
LMF2

LOC100129518
LOC100133286
LOC100288162
LOC100289580
LOC100506100
LOC101448202
LOC101927751
LOC101927765
LOC101928034
LOC102546298
LOC102723692
LOC102724927
LOC284454
LOC644936
LOC730183
LPAR6
LRFN4
LRIG3
LRP3
LRRC45
LXN
MAP1S
MAP3K8
MBD3
MEST
MFAP5
MFSD10
MFSD12
MFSD3
MFSD7
MGLL
MIB2
MIF
MMP14
MMP17
MN1
MNS1
MT2A
MTFP1
MVD
MYOCD
NBEA
NCLN
NDUFB7
NDUFS7
NETO2
NFKBIA
NFKBIZ
NME3
NOG
NOTCH3
NPAS1
NR4A1
NR4A2
NR4A3
NUAK2
NUBP2
OLFM2
OLFM4
ORAI1
OSGIN1
P4HA3
PDE1C
PDE5A
PEAR1
PER2
PHLDA2
PIEZO1
PIM1
PITPNM1
PKMYT1
PLAC8
PLAG1
PLAU
PLCE1
PLCG2
PLD1
PLEC
PLEKHN1
PMAIP1
PODXL
POLD1
POP3
PPP2R3B
PTGDS
PTGIS
PTGS2
PTK2B

PTX3
PYCRL
RABEP2
RARRES1
RASA4B
RASA4CP
RASD2
RCAN2
RCN3
RECQL4
REL
REV3L
RGN
RHOJ
RHOT2
RNF126
RNF150
RPUSD1
SAMD12
SBSN
SBSPON
SCAND1
SCARF2
SCRIB
SDF2L1
SDPR
SEMA3D
SEMA7A
SHOX
SIM1
SIVA1
SLC16A3
SLC17A9
SLC22A17
SLC25A22
SLC2A12
SLC2A6
SLC38A5
SLC52A2
SLC9A3R2
SLX1A
SLX1B
SMTN
SNAI1
SPA17
SPHK1
SPON2
STMN2
SYTL2
TBX4
TCIRG1
TELO2
TINAGL1
TLR4
TMEM132A
TMEM158
TMEM176A
TMSB15A
TMTC2
TNC
TONSL
TOX
TPD52L1
TPGS1
TRPC6
TRPV2
TSPAN13
TTYH3
TUBB3
UBTD1
UCN2
VWA5A
WDR90
WNT5B
XYLT1
ZFHX4
ZNF185
ZNF213
ZNF668
ZNHIT2

**Table S4: Oligonucleotides**

**siRNAs**

| | |
|---|---|
| siRASD1-3 | CCGCGAUGAUCAAGAAGAUdTdT |
| siRASD1-4 | CGACACAACCUAAGGAGGAdTdT |
| siRRAD-2 | CCAUUGUAGUGGACGGAGAdTdT |
| siRRAD-3 | AGGCAUCACUCAUGGUCUAdTdT |

**qPCR primers**

| Name | Sequence | Amplicon length |
|---|---|---|
| CXCL8_fwd | agacagcagagcacacaagc | 62 bp |
| CXCL8_rev | atggttccttccggtggt | |
| GAPDH_fwd | agccacatcgctcagacac | 66 bp |
| GAPDH_rev | gcccaatacgaccaaatcc | |
| GMNN_fwd | gcatctggatctcttgttgga | 126 bp |
| GMNN_rev | ttcactagattctgggacaataacc | |
| IL11_fwd | agctgcaaggtcaagatggt | 61 bp |
| IL11_rev | agattgtttccagtttgctatgg | |
| NQO1_fwd | cggctttgaagaagaaaggat | 110 bp |
| NQO1_rev | cgcagggtccttcagtttac | |
| RASD1_fwd | cctctccatcctcacaggag | 114 bp |
| RASD1_rev | aggcaagacttggtgtcgag | |
| RRAD_fwd | gacgagagcgtttacaaggtg | 129 bp |
| RRAD_rev | ggagcgatcataggtgtgc | |
| TOP2A_fwd | tccacgatacatctttacaatgc | 126 bp |
| TOP2A_rev | aaaaacttcaacgtgtgatcatct | |
| UL29_fwd | ccatcatctcctcgcttagg | 63 bp |
| UL29_rev | agctgcagatcgaggactg | |
| UL6_fwd | aggacggctgggtaaaggt | 60 bp |
| UL6_rev | ggagaatctcgcggaacag | |
